# Identification of neuron-glia signaling feedback in human schizophrenia using patient-derived, mix-and-match forebrain assembloids

**DOI:** 10.1101/2024.12.22.629557

**Authors:** Eunjee Kim, Yunhee Kim, Soojung Hong, Inha Kim, Juhee Lee, Kwanghwan Lee, Myungmo An, Sung-Yon Kim, Sanguk Kim, Kunyoo Shin

**Affiliations:** Institute of Molecular Biology and Genetics, Seoul National University, Seoul 08826, Republic of Korea; School of Biological Sciences, College of Natural Sciences, Seoul National University, Seoul 08826, Republic of Korea; Department of Life Sciences, Pohang University of Science and Technology, Pohang, Gyeongbuk 37673, Republic of Korea; Department of Chemistry, Seoul National University, Seoul 08826, Republic of Korea

**Keywords:** Brain assembloid, brain organoid, human brain development, schizophrenia

## Abstract

Although abnormal activities across multiple cell types are believed to contribute to the development of various neurodevelopmental disorders, current brain organoid technologies fall short in accurately modeling the dynamic cellular interactions in the human brain. Recently, we developed a cellular reconstitution technology to create human forebrain assembloids with enhanced cellular diversity, representing dynamic interactions between neurons and glial cells. Here, we created patient-derived, mix-and-match forebrain assembloids, in which neurons, astrocytes, and microglia from both healthy individuals and schizophrenia patients were reconstituted in a combinatorial manner, and identified aberrant cellular interactions between neurons and glial cells in human schizophrenia. At the early stage, schizophrenia forebrain assembloids showed premature neurogenesis induced by the abnormal proliferation and differentiation of neural progenitor cells. Integrated modular analysis of gene expression in post-mortem schizophrenia brain tissue and brain assembloids found increased expression of tumor protein p53 (TP53) and nuclear factor of activated T-cells 4 (NFATC4), which functioned as master transcriptional regulators to epigenetically reprogram the transcriptome involved in the cellular dynamics of neuronal progenitor cells, leading to premature neurogenesis. At the later stage, we observed weakened structures of laminar organization of the cortical layers in forebrain assembloids and identified the neuron-dependent transcriptional plasticity of glial cells and their altered signaling feedback with neurons, in which neuronal urocortin (UCN) and protein tyrosine phosphatase receptor type F (PTPRF) elicited the expression of Wnt family member 11 (WNT11) and thrombospondin 4 (THBS4) in astrocytes and microglia, respectively. These aberrant signaling axes altered the neuronal transcriptome associated with neuronal response to various stimuli and synthetic processes of biomolecules, resulting in reduced synapse connectivity. Thus, we elucidated developmental stage-specific, multifactorial mechanisms by which dynamic cellular interplay among neural progenitor cells, neurons, and glial cells contribute to the development of the human schizophrenia brain. Our study further demonstrated the potential and applicability of patient-derived forebrain assembloid technology to advance our understanding of the pathogenesis of various neurodevelopmental disorders.

## Introduction

Schizophrenia (SCZ) is a debilitating psychiatric disorder, characterized by positive, negative, and cognitive symptoms, that affects approximately 1% of the worldwide population ^1,2^. Due to the polygenic nature and the diverse risk factors including genetic and pre/postnatal environmental factors, pathophysiology and mechanistic understanding are extremely restricted despite extensive recent research. According to the current neurodevelopmental hypothesis, the pathogenesis of human SCZ is accounted for human brains predisposed to embryonic structural defects, likely caused by genetic factors, and *in utero* exposures to various risk factors such as infections, malnutrition, and exposure to toxins ^3–5^, that undergo the onset of SCZ following exposure to postnatal environmental factors ^6,7^.

Despite the abundance of genomic resources as well as common structural features identified by recent studies, challenges in studying and understanding the pathogenesis of human SCZ still remain. Particularly, one of the major difficulties in understanding the basis of human SCZ stems from the genetic complexity and polygenic nature of this disease. For instance, questions such as how a large number of common variants and risk genes found in patients with SCZ interact with each other at different neurodevelopmental stages of the human brain and what changes in these genetic interactions cause the postnatal onset of human SCZ remain unanswered ^8–11^. This mechanistic understanding requires an experimental model platform that allows for genetic manipulation of the human brain at different stages of neurodevelopment, enabling the dissection of the molecular basis of disease pathogenesis and overcoming the current limitations of using animal models and post-mortem human brains. Several studies have used two-dimensional culture models with various cell types derived from patient-specific induced pluripotent stem cells (iPSCs) ^12–14^. However, these approaches are limited in that they cannot represent the development of the human brain as well as the cellular complexity and dynamic interplay between various cell types, such as neuronal progenitor cells (NPCs), glial cells, and excitatory and inhibitory neurons, in which their alterations may be one of the major causes for the development of human SCZ.

Recently, brain organoid technology has been developed and serves as an experimental platform to model various neurodevelopmental disorders. Brain organoids are three-dimensional (3D), self-organizing tissues derived from pluripotent stem cells, evolving over several years from whole-brain organoids ^15,16^ to region-specific organoids ^17–25^. However, despite their value in studying the human brain, current brain organoids exhibit limited cellular diversity and fail to capture the dynamic cellular interactions found in the human brain. In response, we recently developed a stepwise, module-based cellular reconstitution technology to sequentially build human forebrain assembloids ^26,27^. These assembloids not only demonstrated a high degree of consistency and uniformity with minimal heterogeneity and batch effects, but also exhibited enhanced cellular diversity, representing dynamic interactions between neurons and glial cells. In this study, we created patient-derived, mix-and-match forebrain assembloids by reconstituting neurons, astrocytes, and microglia from both healthy individuals and SCZ patients in various combinations, and identified altered cellular interactions between neurons and glial cells in human SCZ. Our findings thus demonstrate the potential of patient-derived forebrain assembloids in advancing our understanding of the pathogenesis of various human neurodevelopmental disorders, such as SCZ, during human brain development.

## Results

### Aberrant proliferation and differentiation of NPCs in the early developmental stage of SCZ patient-derived forebrain assembloids

As altered gene expressions and abnormal cellular activity of multiple cell types have been hypothesized to contribute to the development of many neurodevelopmental disorders, we tested the applicability of our newly-conceived forebrain assembloids ^27^ to modelling one of the neurodevelopmental disorders, human SCZ, by creating patient-derived forebrain assembloids from human SCZ patients (Fig. 1a-g and Supplementary Fig. 1a-e). We initially obtained patient-specific fibroblasts, B-lymphocytes or iPSCs for eight patients from the NIGMS Human Genetic Cell Repository (Supplementary Data 1). Patient fibroblasts or B-lymphocytes of the eight SCZ patients were then reprogrammed to generate patient-specific iPSCs (Supplementary Fig. 1a) ^28,29^. All eight iPSCs were manipulated to develop into forebrain assembloids specific to patients with SCZ to model the biological and developmental aspects of the human SCZ brain (Supplementary Fig. 1b-e).

**Fig. 1.**
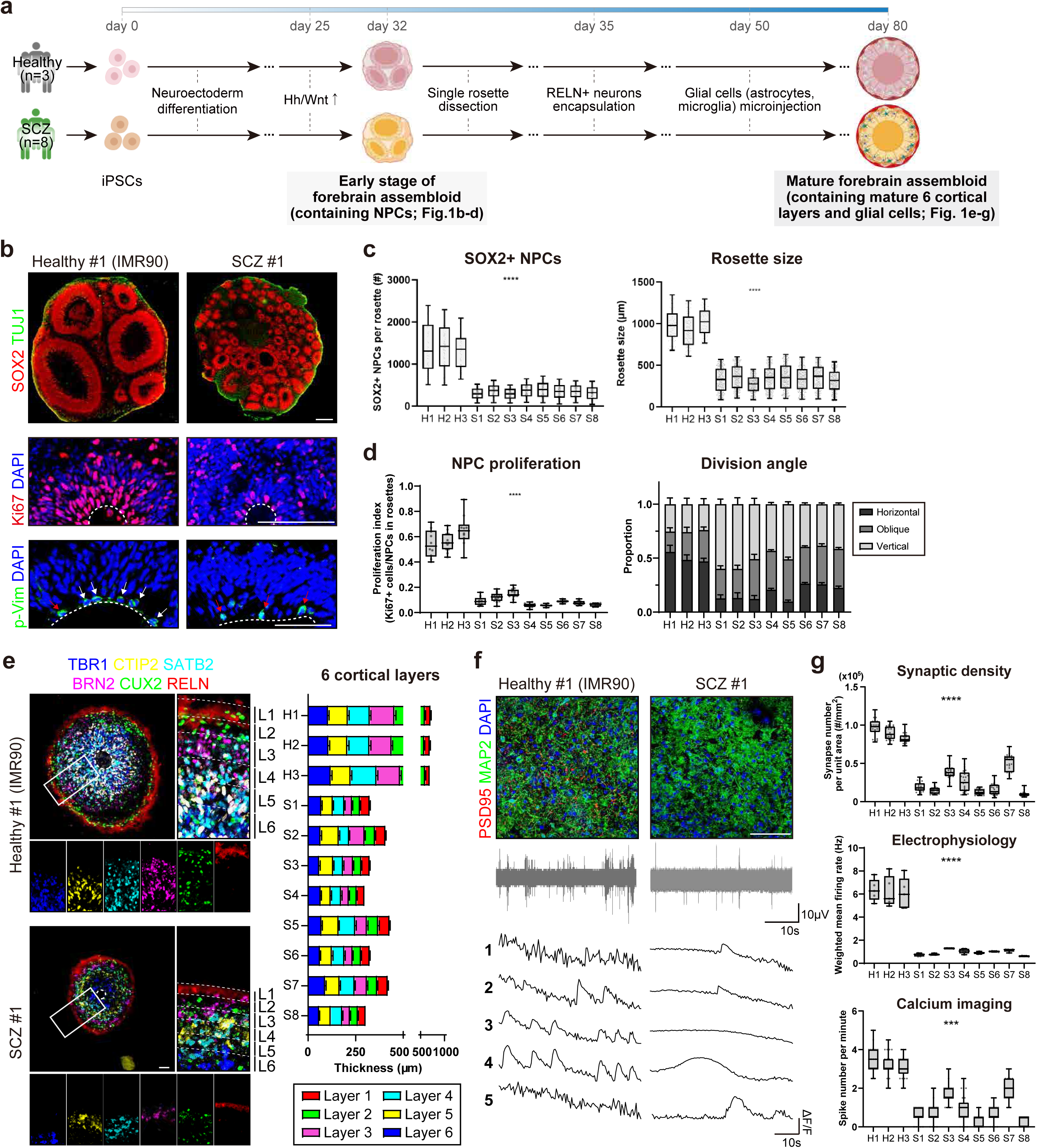
SCZ patient-derived forebrain assembloids show premature neurogenesis and weakened cortex organization with reduced synaptic connectivity. (a) Experimental strategy for the phenotypic analysis of forebrain assembloids at the early stage (d32) and late stage (d80) derived from three healthy individuals and eight SCZ patients. (b) (top) Representative images of early forebrain organoids (d32) immunostained with SOX2 and TUJ1. Scale bars, 100 μm. (middle) Representative images of early brain organoids (d32) immunostained with Ki67. The dotted lines demarcate the border between the VZ and the ventricle. Scale bars, 100 μm. (bottom) Representative images of early brain organoids (d32) immunostained with phospho-vimentin (p-Vim). The dotted lines demarcate the border between the VZ and the ventricle. The white arrows indicate NPCs that underwent horizontal division; the red arrows indicate NPCs that underwent oblique or vertical division. Scale bars, 50 μm. (c) (left) Quantification of the NPC population by counting the number of SOX2-positive cells in each rosette. The number of NPCs in all rosettes was quantified in two sections of one organoid. All organoids in each group are counted. n = 78 (H1), n = 136 (H2), n = 81 (H3), n = 410 (S1), n = 437 (S2), n = 401 (S3), n = 368 (S4), n = 372 (S5), n = 383 (S6), n = 324 (S7), n = 397 (S8). Significance was calculated using a nested *t*-test. Center line, median; whiskers, min to max (show all points). (right) Quantification of the rosette size by measuring the diameter of SOX2+ rosette. All rosettes were quantified in two sections of one organoid. All organoids in each group are counted. n = 78 (H1), n = 136 (H2), n = 81 (H3), n = 410 (S1), n = 437 (S2), n = 401 (S3), n = 368 (S4), n = 372 (S5), n = 383 (S6), n = 324 (S7), n = 397 (S8). Significance was calculated using a nested *t*-test. Center line, median; whiskers, min to max (show all points). (d) (left) Quantification of NPC proliferation by counting Ki67-positive cells as a percentage of the total SOX2-positive cells in each section. NPC proliferation was quantified in two sections of one organoid. All organoids in each group are counted. n = 10 (H1), n = 12 (H2), n = 10 (H3), n = 10 (S1), n =14 (S2), n = 12 (S3), n = 12 (S4), n = 10 (S5), n = 12 (S6), n = 10 (S7), n =10 (S8). Significance was calculated using a nested *t*-test. Center line, median; whiskers, min to max (show all points). (right) The proportion of NPCs with the horizontal, oblique, or vertical orientation of cell division in each section (0–30°: horizontal, 30–60°: oblique, and 60–90°: vertical). The proportion of horizontal, oblique, or vertical division was quantified based on the average division angle of cells in each section. Two sections were quantified for one organoid. All organoids in each group are counted. n = 10 (H1), n = 12 (H2), n = 10 (H3), n = 10 (S1), n =14 (S2), n = 12 (S3), n = 12 (S4), n = 10 (S5), n = 12 (S6), n = 10 (S7), n =10 (S8). (e) (left) Representative images of six cortical layers in healthy and SCZ assembloids (d80). Three serial sections at 8-μm intervals were stained for TBR1/CTIP2, SATB2/RELN, and BRN2/CUX2, respectively. Merged images are shown in the panel on the left. Scale bars, 100 μm. (right) Quantification of the cortical thickness of each of the 6 layers. The six cortical layers were delineated based on the expression of specific markers associated with each layer of the cortex (TBR1 for layer 6, CTIP2 for layer 5, SATB2 for layer 4, BRN2 for layer 3, CUX2 for layer 2, and RELN for layer 1). For layer quantification, the image of cortical plate of forebrain assembloids is evenly divided into 15 bins, spanning from apical to basal directions. For each bin, the proportion of layer-specific marker-positive cells is calculated as [number of layer-specific marker-positive cells / number of total neurons]. Each bin is assigned to one of the six cortical layers based on the major cell population comprising each bin, determined by the proportion of layer-specific marker-positive cells. The thickness of six cortical layers was measured in four independent regions of each section (at 90-degree angle intervals), and two sections were quantified for one assembloid (Five biological replicates were evaluated; n=40). (f) (top) Representative images of healthy and SCZ assembloids (d80) immunostained for neurons (MAP2) and synapses (PSD95). Scale bars, 50 μm. (middle) Representative images of a recording plot from a single electrode of a MEA in healthy and SCZ assembloids. (bottom) Representative images of calcium imaging analysis of selected cells in healthy and SCZ assembloids. (g) (top) Quantification of synaptic density by calculating the number of synapses per unit area (mm^2^) in each section. Five sections were quantified for one assembloid (Five biological replicates were evaluated; n = 25). Significance was calculated using a nested *t*-test. Center line, median; whiskers, min to max (show all points). (middle) Quantification of the weighted mean firing rate in healthy and SCZ assembloids measured by MEA. Recordings were performed for thirty minutes. The signals from all active electrodes were quantified for each assembloid (Five biological replicates were evaluated; n = 5). Significance was calculated using a nested *t*-test. Center line, median; whiskers, min to max (show all points). (bottom) Quantification of spike number per minute analyzed by calcium imaging of selected cells in healthy and SCZ assembloids. The number of spikes and the amplitude were quantified in five selected cells of one assembloid (Five biological replicates were evaluated; n = 25). Significance was calculated using a nested *t*-test. Center line, median; whiskers, min to max (show all points).

Because human SCZ brains are likely to develop through multifactorial mechanisms, which are characterized by various alterations in cellular dynamics and gene expression at different stages during the brain development ^14,30,31^, we examined the forebrain assembloids derived from patients with SCZ at two distinct developmental stages, early and late, to investigate the developmental stage-specific mechanisms (Fig. 1a). Early forebrain organoids are characterized by a high degree of NPC proliferation without distinct cortex specification. Early-stage organoids, derived from all eight SCZ patients, had a reduced number of SOX2-positive cells with decreased proliferation of NPCs, leading to the formation of small rosettes without well-defined ventricles and a thinner VZ (Fig. 1b,c and Supplementary Fig. 2a,b). In addition, defective proliferation of NPCs was accompanied by premature differentiation characterized by increased oblique and vertical cell divisions, with a low rate of horizontal divisions in organoids of these eight SCZ patients (Fig. 1b,d and Supplementary Fig. 2a). These data suggest that the early phenotypic defects in SCZ brains are due to decreased cell proliferation and the premature differentiation of NPCs.

### Weakened laminar organization of cortical layers and reduced synaptic connectivity in the late developmental stage of SCZ patient-derived forebrain assembloids

Next, by reconstituting patient-specific early organoids with RELN-expressing neurons and patient-derived glial cells, we created eight patient-derived SCZ forebrain assembloids to examine structural and functional alterations at the late stage of SCZ brain development (Fig. 1a). Interestingly, all SCZ forebrain assembloids displayed proper laminar organization of the six cortical layers, as seen in healthy forebrain assembloids (Fig. 1e and Supplementary Fig. 2c).

However, a significant reduction in the overall thickness of each layer was seen (Fig. 1e and Supplementary Fig. 2c), which is likely to result from early defects in cell proliferation of NPCs (Fig. 1b,d and Supplementary Fig. 2a). We also found that compared with healthy controls, the functional connectivity in SCZ brain assembloids was dramatically reduced, which is characterized by lower levels of synaptic density, neural excitability, and synapse transmission (Fig. 1f,g and Supplementary Fig. 2d). Taken together, these findings indicate that the human SCZ brain shows weakened laminar organization of cortical layers and impaired synaptic functions at the late stage of neurodevelopment.

### TP53 and NFATC4 reprograms the transcriptome landscape of NPCs to induce premature neurogenesis

Integrative transcriptome analysis with early forebrain organoids and post-mortem brain tissue was performed to investigate the underlying mechanisms of the early defects in NPC proliferation in the SCZ brain (Fig. 2a,b and Supplementary Fig. 3a-d). Initial analysis, combined with gene ontology (GO) analysis, of early brain organoids identified 3,652 differentially expressed genes (DEGs) – 905 of which are associated with neurodevelopment and neuronal differentiation pathways (Supplementary Fig. 3a,b; Supplementary Data 2). Considering the difficulty of relating individual DEGs to NPC proliferation phenotypes due to a large number of DEGs, we performed gene expression module analysis, which required the use of a large-scale transcriptome database from brain tissue samples collected from hundreds of patients with SCZ (PsychENCODE ^9,32^; Fig. 2a). Co-expression module analysis of post-mortem RNA-seq database identified nine SCZ-associated gene expression modules (ME13, 23, 26, 27, 33, 58, 62, 64 and 68) among 73 human development-associated modules (ME1 to ME73) ^33^ (Supplementary Fig. 3c; Supplementary Data 3). A subsequent analysis of the nine SCZ-associated modules for temporal gene expression revealed that three gene expression modules were associated with the early stage of development (ME13, 26 and 58) (Supplementary Fig. 3c). Further analysis of cell-type enrichment found that module 13 (ME13) was enriched in NPCs (Supplementary Fig. 3c). This finding was further supported by GO analysis data, which showed that ME13 is associated with NPC proliferation and differentiation (Fig. 2b). Based on recent studies on the strong association between altered expression of epigenetic regulators and the development of human SCZ ^33^, we attempted to identify potential master regulators that epigenetically control the early transcriptome associated with ME13. A transcription factor (TF) enrichment test revealed twelve candidate genes that can function as master regulators (Fig. 2b; Supplementary Data 3), all of which were also identified in the TF enrichment test for eight SCZ forebrain organoids (Supplementary Fig. 3d; Supplementary Data 2). Further analysis with quantitative RT-PCR (qRT-PCR) finally identified TP53 and NFATC4, whose expression was significantly increased in all eight SCZ forebrain organoids in comparison to healthy organoids (Fig. 2c and Supplementary Fig. 3e).

**Fig. 2.**
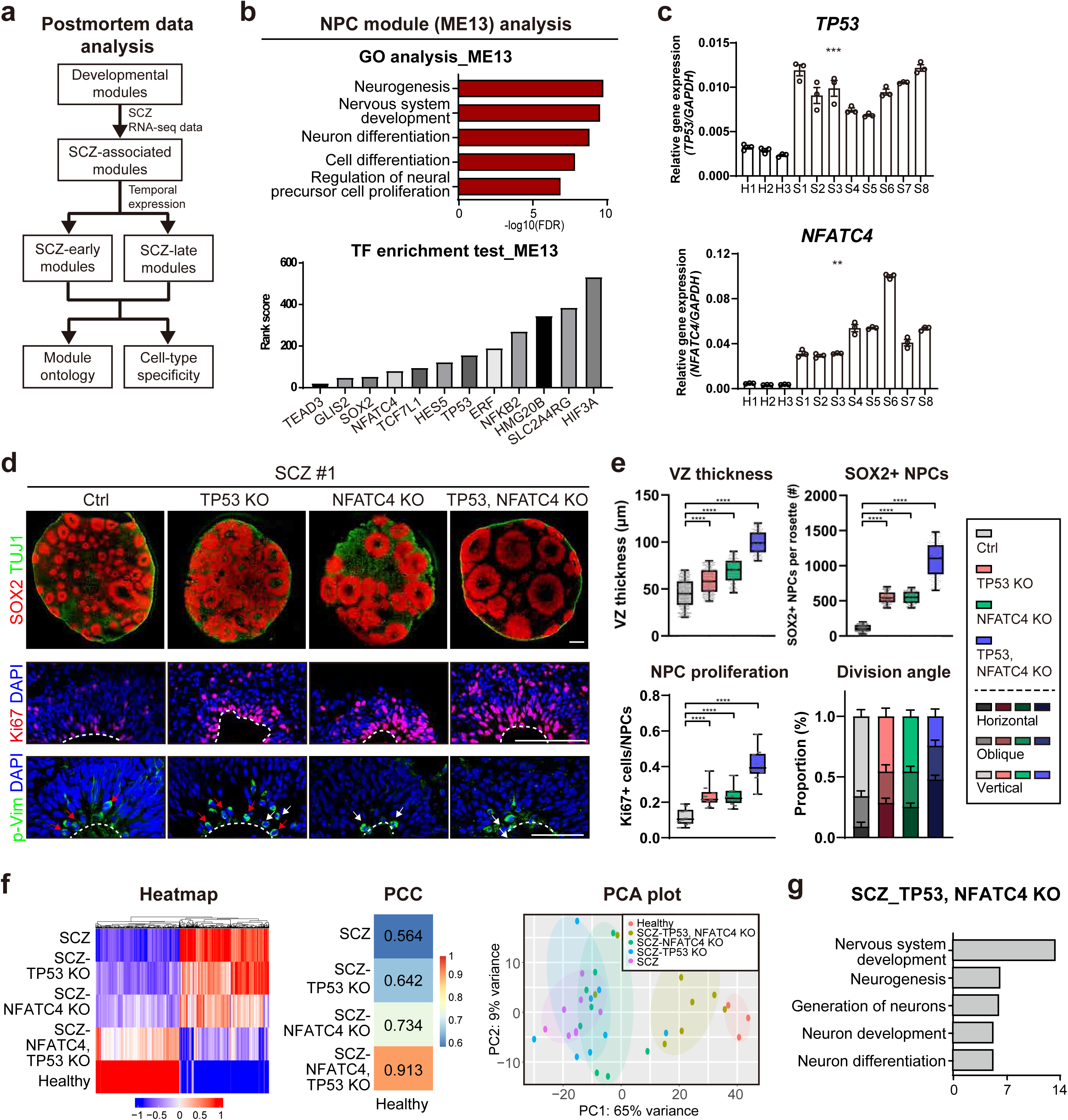
Reprogramming of the transcriptomic landscape in neural progenitor cells by TP53 and NFATC4 at the early stage of human SCZ brain development. (a) Schematic illustration of the co-expression network analysis of bulk RNA-seq data from post-mortem brain tissue. (b) (top) GO analysis of the NPC module (ME13). (bottom) Transcription factor (TF) enrichment analysis for genes within the NPC module (ME13). (c) Relative expressions of *TP53* and *NFATC4* in early forebrain organoids derived from healthy individuals and SCZ patients (Three biological replicates were evaluated; n = 3). Significance was calculated using a nested *t*-test. (d) (top) Representative images of control, only TP53-KO, only NFATC4-KO, and both TP53- and NFATC4-KO organoids derived from a SCZ patient (SCZ #1) immunostained with SOX2 and TUJ1. Scale bars, 100 μm. (middle) Representative images of control, only TP53-KO, only NFATC4-KO, and both TP53- and NFATC4-KO organoids derived from a SCZ patient (SCZ #1) immunostained with Ki67. The dotted lines demarcate the border between the VZ and the ventricle. Scale bars, 100 μm. (bottom) Representative images of control, only TP53-KO, only NFATC4-KO, and both TP53- and NFATC4-KO organoids derived from a SCZ patient (SCZ #1) immunostained with p-vim. The dotted lines demarcate the border between the VZ and the ventricle. The white arrows indicate NPCs that underwent horizontal division; the red arrows indicate NPCs that underwent oblique or vertical division. Scale bars, 50 μm. (e) (top left) Quantification of VZ thickness by measuring the thickness of VZ from two independent regions in each rosette (at 90-degree angle intervals). The VZ thickness of all rosettes was quantified in two sections of one organoid. All organoids in each group are counted. n = 792 (Ctrl), n = 538 (TP53-KO), n = 290 (NFATC4-KO), n = 220 (TP53/NFATC4-KO). Significance was calculated using an unpaired *t*-test. Center line, median; whiskers, min to max (show all points). (top right) Quantification of the NPC population by counting the number of SOX2-positive cells in each rosette. The number of NPCs in all rosettes was quantified in two sections of one organoid. All organoids in each group are counted. n = 396 (Ctrl), n = 269 (TP53-KO), n = 145 (NFATC4-KO), n = 110 (TP53/NFATC4-KO). Significance was calculated using an unpaired *t*-test. Center line, median; whiskers, min to max (show all points). (bottom left) Quantification of NPC proliferation by counting Ki67-positive cells as a percentage of the total SOX2-positive cells in each section. NPC proliferation was quantified in two sections of one organoid. All organoids in each group are counted. n = 10 (Ctrl), n = 10 (TP53-KO), n = 12 (NFATC4-KO), n = 10 (TP53/NFATC4-KO). Significance was calculated using an unpaired *t*-test. Center line, median; whiskers, min to max (show all points). (bottom right) The proportion of NPCs with the horizontal, oblique, or vertical orientation of cell division in each section (0– 30°: horizontal, 30–60°: oblique, and 60–90°: vertical). The proportion of horizontal, oblique, or vertical division was quantified based on the average division angle of cells in each section. Two sections were quantified for one organoid. All organoids in each group are counted. n = 10 (Ctrl), n = 10 (TP53-KO), n = 12 (NFATC4-KO), n = 10 (TP53/NFATC4-KO). (f) RNA-seq analysis of control, TP53-KO, NFATC4-KO, and both TP53- and NFATC4-KO organoids derived from SCZ patients, as well as organoids derived from healthy individuals. (left) Heatmap representation of RNA-seq analysis of indicated forebrain organoids using selected DEGs (log2 fold change >1, p-value < 0.001). (middle) Pearson correlation coefficient obtained by using selected DEGs (log2 fold change >1, p-value < 0.001) of each organoid. (right) PCA plot from RNA-seq analysis of indicated forebrain organoids analyzed by using selected DEGs (log2 fold change >1, p-value < 0.001). (g) GO analysis of TP53- and NFATC4-KO forebrain organoids using selected DEGs (FDR < 0.01).

To obtain direct experimental evidence and mechanistic insights for the roles of TP53 and NFATC4 during the early development of the SCZ brains, we ablated the expression of TP53 and NFATC4 in SCZ forebrain organoids (Supplementary Fig. 4a,b). We found that in the absence of TP53 and NFATC4 expression, the resulting organoids exhibited enhanced proliferation of NPCs, with a concomitant decrease in oblique and vertical divisions, leading to an increased number of SOX2-positive NPCs per rosette (Fig. 2d,e and Supplementary Fig. 4c-e), which is similar to that seen in healthy organoids (Fig. 1b-d and Supplementary Fig. 2a,b).

Furthermore, gene expression analysis revealed that the transcriptome of SCZ forebrain organoids in the absence of TP53 and NFATC4 expression showed a high degree of similarity with the transcriptome of healthy forebrain organoids (Fig. 2f; Supplementary Data 4). GO analysis identified pathways associated with cell proliferation in TP53-ablated forebrain organoids and ones associated with cell differentiation in NFATC4-ablated organoids (Fig. 2g; Supplementary Data 4). These data suggest that the increased expression of TP53 and NFATC4 in SCZ brain organoids induces premature neurogenesis by altering transcriptome landscape associated with cellular proliferation and differentiation.

To further confirm the roles of TP53 and NFATC4 in the development of early SCZ brains, we manipulated healthy forebrain organoids to express TP53 and NFATC4 at high levels (Supplementary Fig. 5a,b). The resulting organoids showed similar phenotypic changes observed in SCZ forebrain organoids, such as smaller rosettes with less SOX2-positive NPCs, a thinner VZ, and a lower degree of cellular proliferation with increased asymmetry of cell division (Supplementary Fig. 5c-e). Moreover, transcriptome analysis showed that the gene expression profile of healthy forebrain organoids overexpressing TP53 and NFATC4 displayed a high similarity with that of SCZ forebrain organoids (Supplementary Fig. 5f; Supplementary Data 5), having altered gene expression associated with cell proliferation and cell differentiation (Supplementary Fig. 5g). Taken together, these findings suggest that TP53 and NFATC4 altered the transcriptomic landscape in the SCZ brain, resulting in the induction of early phenotypic changes through defective proliferation and early differentiation of NPCs.

### Generation of combinatorial, mix-and-match forebrain assembloids

Several studies have suggested a strong association between altered cellular interactions of multiple cell types and the development of human SCZ ^14,30,31^. To elucidate the altered nature of the dynamic interplay between the three major cell types of the human brain – neurons, astrocytes, and microglia – in SCZ brains, we created ‘mix-and-match forebrain assembloids’ by reconstituting patient-derived forebrain organoids with patient-derived glial cells in a combinatorial manner, resulting in eight different forms of forebrain assembloids (Fig. 3a).

**Fig. 3.**
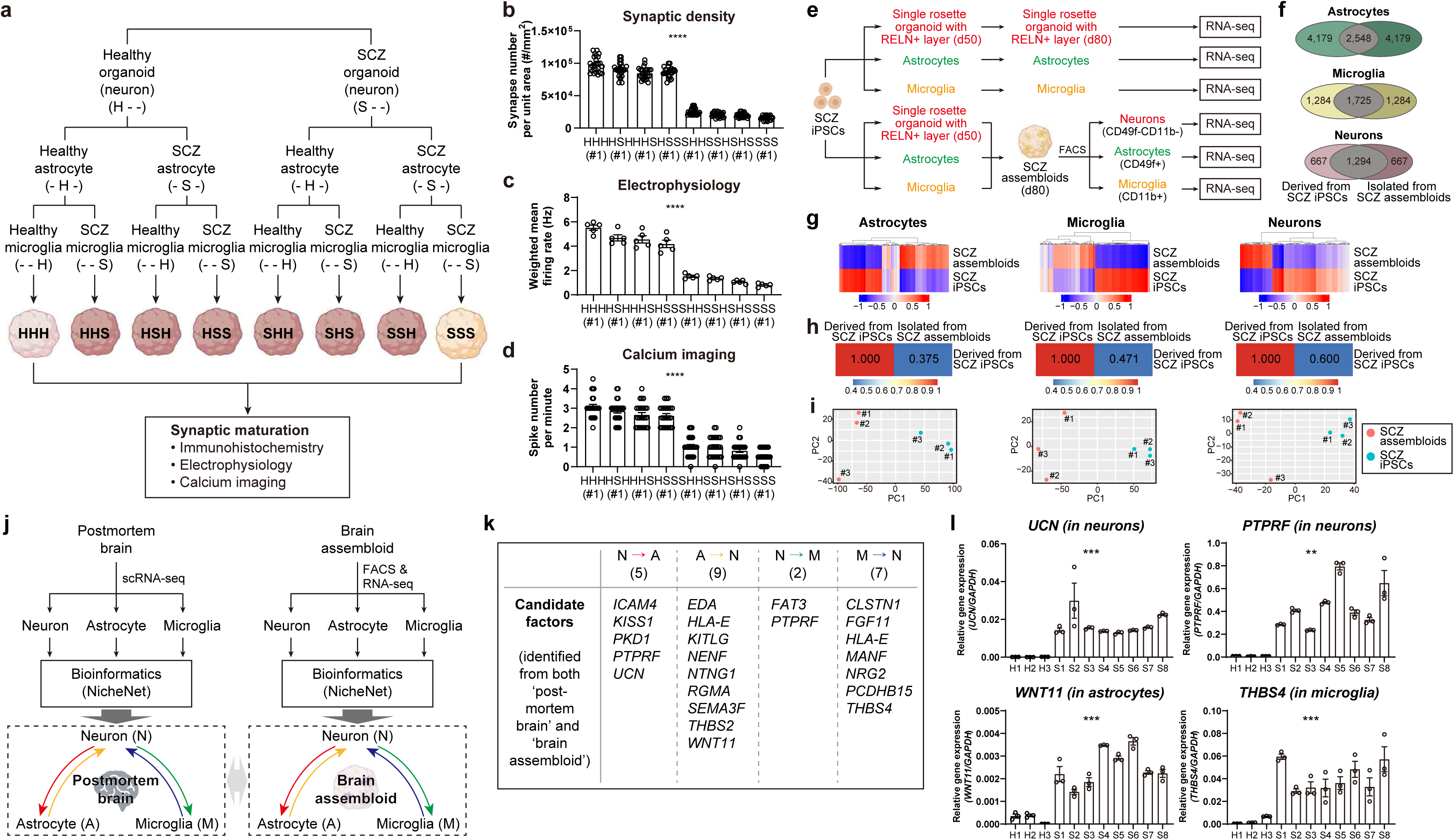
Creation of combinatorial, mix-and-match forebrain assembloids to reveal the neuron-dependent, transcriptional plasticity of glial cells and their alterations at the late stage of human schizophrenia brain development. (a) Experimental scheme of the generation of combinatorial, mix-and-match forebrain assembloids by reconstituting patient-derived forebrain organoids with patient-derived glial cells in a combinatorial manner, resulting in eight different forms of forebrain assembloids. (b) Quantification of synaptic density in the indicated combinatorial assembloids derived from a healthy individual (Healthy #1) and a SCZ patient (SCZ #1) by calculating the number of synapses per unit area (mm^2^) in each section. Five sections were quantified for one assembloid (Five biological replicates were evaluated; n = 25). Significance was calculated using a nested *t*-test. (c) Quantification of the weighted mean firing rate in the indicated combinatorial assembloids derived from a healthy individual (Healthy #1) and a SCZ patient (SCZ #1) measured by MEA. Recordings were performed for thirty minutes. The signals from all active electrodes were quantified for each assembloid (Five biological replicates were evaluated; n = 5). Significance was calculated using a nested *t*-test. (d) Quantification of spike number per minute analyzed by calcium imaging of selected cells in indicated combinatorial assembloids derived from a healthy individual (Healthy #1) and a SCZ patient (SCZ #1). The number of spikes was quantified in five selected cells of one assembloid (Five biological replicates were evaluated; n = 25). Significance was calculated using a nested *t*-test. (e) Schematic diagram of RNA-seq analysis of iPSC-derived and assembloid-derived neurons, astrocytes, and microglia from SCZ patients. Neurons (CD49f-CD11b-), astrocytes (CD49f+ CD11b-), and microglia (CD49f-CD11b+) were isolated from SCZ forebrain assembloids by FACS. (f) Venn diagrams indicating the number of common and differentially expressed genes between SCZ iPSCs-derived and SCZ assembloid-derived cells (astrocytes, microglia, and neurons). (g) Heatmap representations of RNA-seq analysis for astrocytes, microglia, and neurons derived from SCZ iPSCs and SCZ assembloids using selected DEGs (log2 fold change >2, p-value < 0.05). (h) Pearson correlation coefficient obtained using DEGs (log2 fold change >2, p-value < 0.05) in astrocytes, microglia, and neurons derived from SCZ iPSCs and SCZ assembloids. (i) PCA plot of RNA-seq analysis for astrocytes, microglia, and neurons derived from SCZ iPSCs and SCZ assembloids using selected DEGs (log2 fold change >2, p-value < 0.05). (j) Experimental scheme to identify common neuronal and glial factors from the prediction analysis of cell–cell interactions in forebrain assembloids and post-mortem brain tissue. (k) The table displaying neuronal and glial factors identified from the prediction analysis of cell–cell interactions. (l) Relative expressions of *UCN* and *PTPRF* in neurons, and *WNT11* and *THBS4* in astrocytes and microglia, respectively, isolated from healthy and SCZ forebrain assembloids (Three biological replicates were evaluated; n = 3).

Interestingly, functional analysis of the resulting assembloids revealed that, regardless of whether glial cells were derived from healthy or SCZ patients, all four combinatorial forebrain assembloids containing neurons derived from patients with SCZ (SHH, SHS, SSH and SSS) exhibited functional abnormalities, such as decrease of synaptic density, neural excitability, and synapse transmission (Fig. 3b-d and Supplementary Fig. 6a,b), leading to the possibility of the lack of involvement of glial cells in the development of SCZ. To test this possibility, we examined normal and SCZ assembloids in the presence or absence of glial cells and performed comparative functional analysis (Supplementary Fig. 6c). Unexpectedly, we found that healthy forebrain assembloids displayed significantly enhanced synaptic functions in the presence of healthy glial cells (Supplementary Fig. 6d,e), whereas SCZ brain assembloids showed dramatically reduced synaptic functions in the presence of SCZ glial cells (Supplementary Fig. 6d,e). In addition, in the absence of glial cells, healthy forebrain assembloids exhibited reduced functional connectivity, as shown in SCZ brain assembloids (Supplementary Fig. 6d,e). These results clearly demonstrated that—opposite to our initial hypothesis—glial cells played critical roles in the development of SCZ brains with their plasticity. This plasticity of glial cells is likely to be dictated by SCZ neurons, which accounts for the functional abnormalities and other phenotypic changes found in mix-and-match SCZ forebrain assembloids reconstituted with glial cells derived from healthy individuals.

### Altered transcriptomes of neurons and glial cells are mutually dependent in human SCZ

To investigate the plastic nature of glial cells, which might be regulated by their interactions with neuronal cells, we performed a comparative RNA-seq analysis of glial cells derived from SCZ iPSCs before and after being reconstituted to create SCZ forebrain assembloids (Fig. 3e and Supplementary Fig. 6f). Interestingly, the transcriptome and principal component analysis showed that the transcriptomic landscape of glial cells, including astrocytes and microglia, isolated from SCZ assembloids was significantly different from that of the same glial cells before the reconstitution with SCZ neurons (Fig. 3f-i). These results indicate the dynamic nature of glial cells, whose transcriptome landscape is plastic, depending on their interactions with neuronal cells in the human SCZ brain. Notably, SCZ neurons also showed transcriptomic alterations in genes involved in the “detection of stimulus” and “biosynthetic process” based on the GO analysis, after reconstitution with glial cells derived from SCZ iPSCs (Fig. 3f-i and Supplementary Fig. 6g; Supplementary Data 6). These findings demonstrate the transcriptional plasticity of neuronal cells and their dependency on glial cells in SCZ brains.

Taken together, these results suggest the mutual dependency between neurons and glial cells for the development of SCZ brains, and indicate towards the presence of a reciprocal signaling feedback, in which alterations may cause synaptic dysfunction in the late stage of human brain development.

### The heightened activity of the UCN/PTPRF-WNT11/THBS4 signaling axis between neurons and glial cells leads to late developmental defects in the SCZ forebrain assembloids

To elucidate the nature of altered reciprocal signaling between neurons, astrocytes, and microglia, we reasoned that the altered expression of neuronal factors elicited cellular response in glial cells, leading to the increased expression of glial factors, which in turn induced neuronal response to develop synaptic dysfunctions in SCZ brains. To first identify neuronal factors that might induce transcriptional changes in glial cells, we performed computational analysis ^34^ (Fig. 3j and Supplementary Fig. 7a) and identified 223 genes (112 and 111 genes responsible for transcriptome changes in astrocytes and microglia, respectively) among DEGs in iPSC-derived SCZ neurons that coded secretory or junctional proteins and showed a high probability of inducing transcriptome changes in astrocytes and microglia of SCZ forebrain assembloids (Supplementary Fig. 7a; Supplementary Data 7). Among these genes, we selected the top 100 neuronal candidate genes responsible for transcriptome changes in astrocytes and microglia (50 genes each) with the highest regulatory potential scores for further analysis (Supplementary Data 7). In subsequent analysis, we identified a total of 366 genes (161 for astrocytes and 205 for microglia) from the altered transcriptome of astrocytes and microglia derived from SCZ assembloids that can elicit additional transcriptome alterations in neurons of SCZ assembloids. We then selected the top 100 glial candidate genes, 50 each for astrocytes and microglia, with the highest regulatory potential scores for further analysis (Supplementary Data 7). These analyses identified numerous reciprocal signaling circuits between neurons and glial cells, whose activities are likely to be heightened in the human SCZ brain (Supplementary Data 7).

Similar but independent computational analyses using a single-cell RNA-seq database from post-mortem tissues ^35^ (see the Methods section for details) were performed to test whether the consensus of the potential signaling feedback circuits can be obtained (Supplementary Data 8). We first identified 164 neuronal genes (85 and 79 genes responsible for transcriptome changes in astrocytes and microglia, respectively) and 502 glial genes (331 for astrocytes and 171 for microglia). We then selected the top 100 neuronal candidate genes (50 genes each responsible for transcriptome changes in astrocytes and microglia) as well as 100 glial candidate genes (50 each for astrocytes and microglia) according to the regulatory potential scores. We then performed a comparative analysis between brain assembloids and post-mortem tissues and identified several common signaling circuits consisting of five neuronal factors (ICAM4, KISS1, PKD1, PTPRF, and UCN) and nine astrocyte factors (EDA, HLA-E, KITLG, NENF, NTNG1, RGMA, SEMA3F, THBS2, and WNT11) for neuron-astrocyte interaction; two neuronal factors (FAT3 and PTPRF) and seven microglial factors (CLSTN1, FGF11, HLA-E, MANF, NRG2, PCDHB15, and THBS4) for the neuron–microglia interaction (Fig. 3k). Finally, we performed quantitative RT-PCR to confirm the expression of the candidate genes in neurons and glial cells isolated from patient-derived forebrain assembloids (Fig. 3l and Supplementary Fig. 7b-d) and identified two common reciprocal signaling axes, one between the neuron and astrocyte (UCN– WNT11) and the other between neurons and microglia (PTPRF–THBS4) (Supplementary Fig. 7e).

We further confirmed the signaling axes identified by genetically manipulating forebrain assembloids combinatorially (Fig. 4a-m and Supplementary Fig. 8a-n). In forebrain assembloids created by reconstituting UCN/PTPRF-ablated SCZ neurons with SCZ glial cells (Fig. 4a), the expression of WNT11 and THBS4 in astrocytes and microglia, respectively, was significantly decreased and functional connectivity was significantly enhanced (Fig. 4b,c and Supplementary Fig. 8a-d). In forebrain assembloids created by reconstituting SCZ neurons with WNT11-ablated astrocytes and THBS4-ablated microglia (Fig. 4d), the transcriptome alterations associated with the “detection of stimulus” and “biosynthetic process” found in the neurons of SCZ assembloids, were not observed and synaptic functions were restored (Fig. 4e,f and Supplementary Fig. 8e-g). Furthermore, the rescue in functional connectivity of forebrain assembloids with UCN/PTPRF-ablated SCZ neurons and SCZ glial cells was reversed in forebrain assembloids with UCN/PTPRF-ablated SCZ neurons and WNT11- and THBS4-overexpressing astrocytes and microglia, respectively, further confirming the reciprocal signaling between neurons and glial cells (Fig. 4g-i and Supplementary Fig. 8h-j). Lastly, to differentiate the role of astrocytes and microglia, we created two different forebrain assembloids by reconstituting SCZ neurons with WNT11-ablated astrocytes and SCZ microglia (Fig. 4j), or with SCZ astrocytes and THBS4-ablated microglia (Fig. 4l). Although both assembloids showed a similar degree of partial restoration of synaptic functions (Supplementary Fig. 8k-n), SCZ neurons derived from assembloids with WNT11-ablated astrocytes and with THBS4-ablated microglia showed distinct transcriptional alterations in that altered transcriptome in assembloids with WNT11-ablated astrocytes was associated with only the “biosynthetic process” whereas that in assembloids with THBS4-ablated microglia is enriched in only the “detection of stimulus” (Fig. 4k,m). These results suggest two distinct neuronal alterations in the regulation of neuronal response to various stimuli and synthetic processes of biomolecules induced by astrocyte and microglia, respectively, for the development of late SCZ brains.

**Fig. 4.**
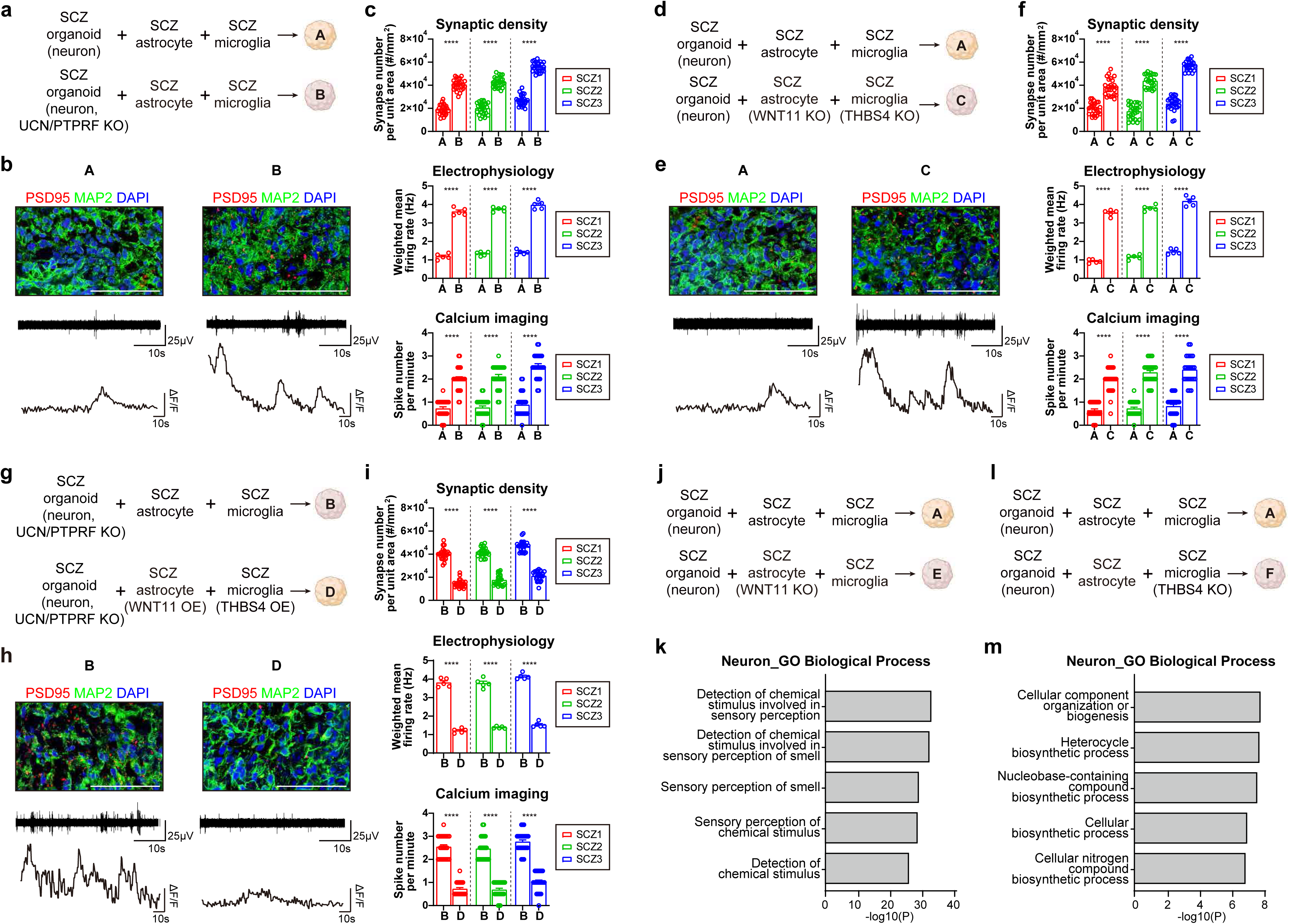
The heightened activity of the UCN/PTPRF-WNT11/THBS4 signaling axis between neurons and glial cells leads to the late developmental defects in the SCZ brain. (a, d, g) Experimental scheme of identifying reciprocal signaling between neurons and glial cells using genetically manipulated forebrain assembloids. (b, e, h) (top) Immunostaining analysis of synaptic density in indicated forebrain assembloids derived from a SCZ patient (SCZ #1). Scale bars, 50 μm. (middle) Representative images of a recording plot from a single electrode of a MEA in indicated forebrain assembloids derived from a SCZ patient (SCZ #1). (bottom) Representative images of calcium imaging analysis of selected cells in indicated forebrain assembloids derived from a SCZ patient (SCZ #1). (c, f, i) Quantification of synaptic density, weighted mean firing rate, and spike number per minute of genetically manipulated forebrain assembloids derived from SCZ patients (SCZ #1, 2, 3). (top) Quantification of the synaptic density in indicated forebrain assembloids by calculating the number of synapses per unit area (mm^2^) in each section. Five sections were quantified for one assembloid (Five biological replicates were evaluated; n = 25). Significance was calculated using an unpaired *t*-test. (middle) Quantification of the weighted mean firing rate in the indicated forebrain assembloids measured by MEA. Recordings were performed for thirty minutes. The signals from all active electrodes were quantified for each assembloid (Five biological replicates were evaluated; n = 5). Significance was calculated using an unpaired *t*-test. (bottom) Quantification of spike number per minute analyzed by calcium imaging of selected cells in indicated forebrain assembloids. The number of spikes was quantified in five selected cells of one assembloid (Five biological replicates were evaluated; n = 25). Significance was calculated using an unpaired *t*-test. (j, l) Experimental scheme of the investigation of the role of astrocytes and microglia in inducing SCZ phenotypes at late stages by reconstituting SCZ forebrain assembloids with genetically manipulated glial cells. (k, m) GO analysis of DEGs between neurons isolated from genetically manipulated forebrain assembloids as indicated in panel ‘j’ (k) and ‘l’ (m).

Taken together, our findings suggest that the heightened activity of reciprocal signaling between SCZ neurons and glial cells, which is mediated through the UCN/PTPRF-WNT11/THBS4 signaling axis, induces late phenotypic alterations, such as the decreased functional connectivity through aberrant neuronal responses to various stimuli and defected synthetic processes of biomolecules, in human SCZ brains.

### Restoration of structural, functional, and behavioral abnormalities in chemically-induced mouse model of human SCZ with *in utero* manipulations of master regulators

Finally, to further validate our findings and *in vivo* relevance, we performed animal experiments and behavior analyses using chemically induced mouse model of SCZ (E13-methylazoxymethanol (MAM)-exposed mouse model), which mimics transcriptomic and histological changes as well as neural dysfunctions found in human SCZ patients ^36,37^ (Supplementary Fig. 9a-d). We found that in the brains of chemically-induced SCZ animals, expressions of the early (Tp53 and Nfatc4) and late (Ucn, Wnt11, Ptprf, and Thbs4) genes were increased at the early and late stages of development, respectively, which was consistent with our finding in the assembloid model (Supplementary Fig. 9e-g). Furthermore, these animals showed decreased number of NPCs, reduced cortical thickness, and strong behavior abnormalities which have been known to be associated with human SCZ (Supplementary Fig. 9h-p). Interestingly, *in utero* genetic rescues by suppressing the expressions of early and late genes resulted in the restoration of NPC proliferation and cortical thickness in embryonic and postnatal brains, respectively, of the chemically-induced SCZ animals (Fig. 5a-c and Supplementary Fig. 10a-g), as well as the alleviation of behavioral abnormalities evaluated through open field tests, elevated plus maze tests, and sucrose preference tests (Fig. 5d-h). Notably, animals of which the expressions of only the early genes (Tp53 and Nfatc4) were knocked down showed restored cortical thickness, but only partial restoration in functional connectivity analyzed by behavior tests, which might be due to high levels of pathogenic expressions of the late genes (Fig. 5i-m and Supplementary Fig. 10h-k). Similarly, the suppression of only the late genes (Ucn, Wnt11, Ptprf, and Thbs4) also resulted in a partial restoration in functionality, likely because of the remaining structural defects in the cortical layers caused by pathogenic expressions of the early genes (Fig. 5i-m and Supplementary Fig. 10h-k). These results imply the temporally and functionally distinct roles of these two gene sets as well as the effects of early structural defects on later functional connectivity. Together with our findings in *in vitro* assembloid models, these results strongly suggest the effects of the early expression of TP53 and NFATC4 on the defective formation of early cortical layers as well as the pathological role of the UCN/PTPRF-WNT11/THBS4 signaling axis in the establishment of functional connectivity.

**Fig. 5.**
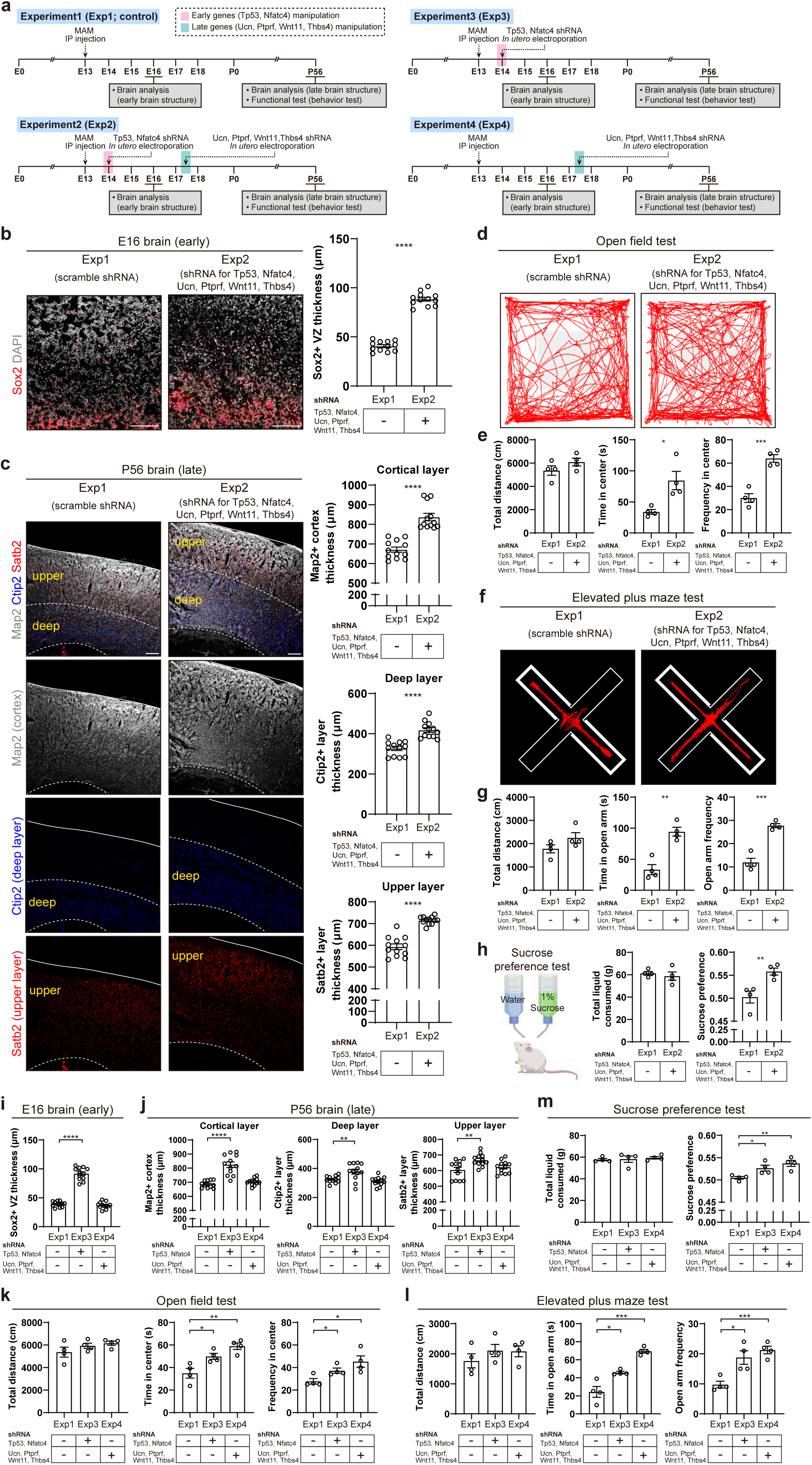
*In utero* genetic manipulation of 6 master regulators restores cortex structure and alleviates behavior abnormalities in the MAM-induced SCZ mouse model. (a) Experimental timeline of gene manipulation, brain analysis, and behavior tests. MAM was administered to pregnant C57BL/6 female mice by intraperitoneal injection, and *in utero* electroporation was performed at E14 and/or E17.5. Electroporated mouse brains were analyzed at E16 or P56 (8 weeks), and mice were analyzed by behavior tests. (b) (left) Representative images of E16 MAM-treated mouse brain cortex electroporated with DNA containing scramble (Exp1) or Tp53/Nfatc4/Ucn/Ptprf/Wnt11/Thbs4 shRNA (Exp2). Coronal sections were immunostained with Sox2. Scale bars, 100 μm. (right) Quantification of the thickness of Sox2+ VZ of E16 MAM-treated mouse brain cortex electroporated with DNA containing scramble (Exp1) or Tp53/Nfatc4/Ucn/Ptprf/Wnt11/Thbs4 (Exp2) shRNA. Three sections per mouse and four mice per group were evaluated (n = 12). Significance was calculated using an unpaired *t*-test. (c) (left) Representative images of P56 MAM-treated mouse brain cortex electroporated with DNA containing scramble (Exp1) or Tp53/Nfatc4/Ucn/Ptprf/Wnt11/Thbs4 shRNA (Exp2). Coronal sections were immunostained with Map2, Satb2, and Ctip2. Scale bars, 100 μm. (right) Quantification of the thickness of Map2+ cortical layer (top), Ctip2+ deep layer (middle), and Satb2+ upper layer (bottom) of P56 MAM-treated mouse brain cortex electroporated with DNA containing scramble (Exp1) or Tp53/Nfatc4/Ucn/Ptprf/Wnt11/Thbs4 shRNA (Exp2). Three sections per mouse and four mice per group were evaluated (n = 12). Significance was calculated using an unpaired *t*-test. (d) Representative track images of open field tests. (e) Quantification of the total distance traveled (left), time spent in center area (middle), and frequency entering center area (right) of P56 MAM-treated mice electroporated with DNA containing scramble (Exp1) or Tp53/Nfatc4/Ucn/Ptprf/Wnt11/Thbs4 shRNA (Exp2). Four mice per group were evaluated (n = 4). Significance was calculated using an unpaired *t*-test. (f) Representative track images of elevated plus maze tests. (g) Quantification of the total distance traveled (left), time spent in open arm (middle), and frequency entering open arm (right) of P56 MAM-treated mice electroporated with DNA containing scramble (Exp1) or Tp53/Nfatc4/Ucn/Ptprf/Wnt11/Thbs4 shRNA (Exp2). Four mice per group were evaluated (n = 4). Significance was calculated using an unpaired *t*-test. (h) Quantification of the total amount of liquid consumed (left) and the sucrose preference (right) of P56 MAM-treated mice electroporated with DNA containing scramble (Exp1) or Tp53/Nfatc4/Ucn/Ptprf/Wnt11/Thbs4 shRNA (Exp2). Four mice per group were evaluated (n = 4). Significance was calculated using an unpaired *t*-test. (i) Quantification of the thickness of Sox2+ VZ of E16 MAM-treated mouse brain cortex electroporated with DNA containing scramble (Exp1), Tp53/Nfatc4 shRNA (Exp3) or Ucn/Ptprf/Wnt11/Thbs4 shRNA (Exp4). Three sections per mouse and four mice per group were evaluated (n = 12). Significance was calculated using an unpaired *t*-test. (j) Quantification of the thickness of Map2+ cortical layer (left), Ctip2+ deep layer (middle), and Satb2+ upper layer (right) of P56 MAM-treated mouse brain cortex electroporated with DNA containing scramble (Exp1), Tp53/Nfatc4 shRNA (Exp3) or Ucn/Ptprf/Wnt11/Thbs4 shRNA (Exp4). Three sections per mouse and four mice per group were evaluated (n = 12). Significance was calculated using an unpaired *t*-test. (k) Quantification of the total distance traveled (left), time spent in center area (middle), and frequency entering center area (right) of P56 MAM-treated mice electroporated with DNA containing scramble (Exp1), Tp53/Nfatc4 shRNA (Exp3) or Ucn/Ptprf/Wnt11/Thbs4 shRNA (Exp4). Four mice per group were evaluated (n = 4). Significance was calculated using an unpaired *t*-test. (l) Quantification of the total distance traveled (left), time spent in open arm (middle), and frequency entering open arm (right) of P56 MAM-treated mice electroporated with DNA containing scramble (Exp1), Tp53/Nfatc4 shRNA (Exp3) or Ucn/Ptprf/Wnt11/Thbs4 shRNA (Exp4). Four mice per group were evaluated (n = 4). Significance was calculated using an unpaired *t*-test. (m) Quantification of the total amount of liquid consumed (left) and the sucrose preference (right) of P56 MAM-treated mice electroporated with DNA containing scramble (Exp1), Tp53/Nfatc4 shRNA (Exp3) or Ucn/Ptprf/Wnt11/Thbs4 shRNA (Exp4). Four mice per group were evaluated (n = 4). Significance was calculated using an unpaired *t*-test.

## Discussion

SCZ is a debilitating psychiatric disorder, characterized by positive, negative, and cognitive symptoms, that affects approximately 1% of the worldwide population ^1,2^. Due to its polygenic nature and the diverse risk factors, including genetic and pre/postnatal environmental influences, the pathophysiology of human SCZ remains poorly understood, despite extensive recent research. According to the current neurodevelopmental hypothesis, the pathogenesis of human SCZ is accounted for human brains, predisposed to embryonic structural defects, that undergo the onset of SCZ following exposure to postnatal environmental factors ^6,7^. To gain a mechanistic understanding of human SCZ development, several studies have used two-dimensional culture models with various cell types derived from patient-specific iPSCs ^12–14^.

However, these approaches are limited in that they cannot represent the development of the human brain as well as the cellular complexity and dynamic interplay between various cell types, such as neuronal progenitor cells, glial cells and excitatory and inhibitory neurons, in which their alterations may be one of the major causes for the development of human neurodevelopmental disorders. In our study, to overcome the current limitations of using iPSC technology, as well as animal and post-mortem brains, we developed patient-derived forebrain assembloids, where the human brain can be genetically manipulated at different stages of neurodevelopment to dissect the molecular basis of disease pathogenesis.

According to our “two-step, multifactorial model”, human SCZ brains develop structural and functional defects at two distinct development stages, and these defects are caused by premature neurogenesis as well as altered cellular interplay between neurons, astrocytes, and microglia (Supplementary Fig. 10l). At the early stage of brain development, increased expression of TP53, known to regulate gene expressions involved in cell cycle arrest and cell survival in a various contexts including the regulation of neurogenesis in the human brain ^38–40^, and NFATC4, which regulates gene expression for the self-renewal and differentiation of progenitors during brain development ^41–43^, reduced the proliferation of NPCs with increased premature differentiation by epigenetically reprogramming the transcriptome involved in the cellular dynamics of NPCs.

At the late stage of brain development, SCZ patient-derived mix-and-match forebrain assembloids revealed altered cellular interactions between neurons and glial cells, mediated through the UCN–WNT11 and PTPRF–THBS4 signaling axes. Notably, the increased expression of all four genes was associated with synaptic dysfunctions according to the analysis of the post-mortem database ^44–47^ (Supplementary Data 8) and showed a strong correlation with cellular communications by functioning as secretory or junctional molecules ^48,49^. UCN, originally identified in the rat midbrain as a member of corticotropin-releasing factor family ^50^, is a secreting protein that is widely expressed within central nervous systems and peripheral tissues in humans. Besides its well-established roles in the control of normal physiology and stress response, the previous study showed that UCN modulated neuronal apoptosis through the regulation of paracrine signalling between dopaminergic neurons and glial cells ^48^, suggesting its additional role as a signalling molecule to communicate with multiple cell types in the brain.

PTPRF, which was identified as another neuronal factors to modulate the interaction with microglia in our study, is a transmembrane protein that belongs to a member of the protein tyrosine phosphatase family. PTPRF is known to be involved in the regulation of cell–cell adhesion, cytoskeleton remodelling and synapse maturation through its interaction with extracellular matrix and neighbouring cells. Recent reports demonstrated that the extracellular subunit (E-subunit) of PTPRF is cleaved to form the secreted fragments, functioning as a signalling ligand to facilitate nerite outgrowth non-cell autonomously ^49,51^. These studies support our proposed mechanisms in which PTPRF, serving as a signalling molecule, mediates cell–cell interaction between neurons and glial cells.

WNT11 and THBS4, based on our integrated computational analysis of glial cells, are the downstream targets in signalling pathways activated by extracellular neuronal UCN and PTPRF, respectively (Supplementary Data 7,8). WNT11, a member of the WNT family of proteins, is known to signal through the non-canonical Wnt pathway. Previous studies in mature neurons suggested that the dysregulated non-canonical pathway induced by WNT11 is associated with various synaptic dysfunctions, such as defected release of neurotransmitters and disorganization of synaptic assembly. Interesting, one study showed that WNT11 was able to induce the changes in gene transcription, which is involved in the detection of mechanical stimulus. These previous findings were consistent with our proposed role of glial WNT11 in inducing transcriptome changes in SCZ neurons, which is associated with stimulus detection, leading to the synapse dysfunction.

THBS4, a member of the thrombospondin protein family, is a glycoprotein that can be secreted to form an extracellular protein and to act as a signalling molecule, depending on the cellular context. Previous studies demonstrated that THBS4 is involved in the formation of functional synapses by mediating cell adhesion, cell-matrix interactions and inflammatory response. THBS4 is believed to function as a secreted molecule that modulates transcription by interacting with the cellular integrins of neighbouring cells and regulating ECM synthesis. These observations are consistent with our findings that in glial cells, THBS4 induces changes in neuronal transcription associated with biosynthetic processes that lead to a dysfunctional synapse. We speculate that the association of WNT11 and THBS4 with the dysfunctional synapse is mediated, in part, by dysregulating cellular interactions between glial cells and neurons and by altering neuronal gene expression.

Taken together, we propose pathogenic mechanisms in which the increased expression of WNT11 and THBS4 in glial cells, induced by neuronal UCN and PTPRF, leads to the formation of a dysfunctional synapse by mediating glial interactions with SCZ neurons which subsequently alters neuronal transcriptomes associated with neuronal response to various stimuli and synthesis of biomolecules. We argue that the identification of surprising plasticity of neurons and glial cells depending on the cellular interactions between each other and the unexpected roles of glial cells in reprogramming neuronal transcriptome to affect the synapse functionality suggests a new conceptual paradigm of how we should understand this complex disease. Furthermore, we strongly believe that our efforts to elucidate signaling feedback between multiple cell types in the human brain is particularly important in that such signaling feedback may lie at the heart of the complex interplay between neurons and glial cells for the human brain development, which is essential to our understanding of various neurodevelopmental disorders in humans.

Another important conceptual advance suggested in this study is that we were able to suggest common mechanisms for the development of human SCZ. Although recent studies revealed a large number of risk variants and thousands of potentially associated genes ^52^ and common structural and functional defects in most human SCZ brains ^53–57^, understanding the common mechanisms of the pathogenesis of human SCZ has been hindered by the genetic complexity and polygenic nature of this disorder. By analyzing gene expression networks in patient-derived SCZ assembloids, our study identified common epigenetic mechanisms through which altered expression of six master regulators mediates cellular behaviors and signaling crosstalk between multiple cell types during human SCZ brain development.

Lastly, this study suggested a new technology to overcome one of the major issues in current large-scale transcriptome analysis for the human brain. Considering that the transcriptomes of human brain are dynamic and flexible depending on different developmental stages and the postnatal status of the human brain samples, most transcriptome databases currently available for human SCZ research are derived from post-mortem brain tissues ^32,47^, which renders the overall transcriptome landscape a snapshot of this particular stage of the human brains. This makes it extremely difficult to investigate dynamic changes in gene expression networks at the different stages of developing human SCZ brains, which are likely to play a major role in the pathogenesis of human SCZ according to a neurodevelopmental hypothesis in which alterations in various cellular signaling pathways during brain development causes defected connectivity of neural circuits, leading to the onset of SCZ in later critical maturational periods. Moreover, an additional layer of difficulty in understanding the biological mechanisms of human SCZ development lies in that human SCZ is likely to result from the alterations in cellular complexity and dynamic interactions between diverse cell types, including neuronal progenitor cells, glial cells, and various types of neurons at different developmental stages of the human brain. Thus, it is imperative to develop human brain models that represent the overall transcriptome of the human SCZ brain at different stages of neurodevelopment in different cell types and to develop experimental model platforms to recapitulate the cellular complexity and dynamic interplay between diverse cell types in the human brains. In this study, we developed SCZ-specific brain assembloids that, we believe, may overcome many limitations in studying the pathogenesis of human SCZ, such as the lack of systems representing different stages of developing human brains with cellular complexity. Combining the integrated modular analysis of transcriptomes with our newly conceived brain assembloids, in which the independent roles of various cell types and their interactions can be investigated through combinatorial reconstitution of patient-derived brain cells into a single organoid, we propose a “two-step, multifactorial mechanism” by which human SCZ brains are developed through premature neurogenesis at an early stage, leading to the weakened structure of laminar organization of cortical layers, and aberrant cellular interactions between neurons and glial cells at later developmental stages, causing reduced synapse connectivity (Supplementary Fig. 10l).

Taken together, our study, which has the potential to profoundly affect our understanding of the role of complex cellular interactions in the development of various brain disorders, will facilitate the establishment of an innovative model platform to study a range of human neurological diseases, whose understanding of pathogenesis requires an organoid system that is capable of representing mature characteristics of functional human brains, including dynamic interplay between neurons and other cell types and will further provide a unique tool for the development of new therapeutic options that can be customized for individual patients.

## Methods

### hiPSC culture

All hiPSCs were maintained on mitomycin C-treated mouse embryonic fibroblasts (MEFs) in hiPSC medium, containing DMEM/F12 (Gibco) supplemented with 20% KnockOut Serum Replacement (Gibco), 1× Glutamax (Gibco), 1× Non-essential amino acids (Gibco), 1% penicillin–streptomycin, 100 μM 2-Mercaptoethanol (Sigma) and 10 ng/ml human bFGF (Peprotech). All hiPSC lines were cultured on each well of a 24-well plate with 0.5 ml of culture media (with 5 x 10^4^ feeder cells per well). The cells were fed daily and passaged by manual dissection at 70% confluence (10 manually-dissected, small colonies of hiPSCs were plated on each well of a 24-well plate). Cells used in this study were negative for mycoplasma contamination (e-Myco Mycoplasma PCR detection kit).

### Generation of patient-derived hiPSCs

hiPSCs (IMR90, female) from the healthy individual were obtained from WiCell. hiPSCs (GM25256, male; GM23338, male) from healthy individuals, hiPSCs (GM23762, male), fibroblasts (GM01835, female; GM02038, male; GM02503, female; GM01792, male) and B-lymphocytes (GM23836, male; GM01487, female; GM01489, male) from patients with SCZ were obtained from the National Institute of General Medical Sciences Cell Repository through the Coriell Institute for Medical Research. SCZ fibroblasts and B-lymphocytes were reprogrammed into iPSCs by retroviral transduction using the Yamanaka factors (Oct4, Klf4, Sox2 and c-Myc) as previously described ^27,58^. Briefly, 1 × 10^5^ human fibroblasts or B-lymphocytes were transduced with retroviruses containing OCT3/4, SOX2, KLF4 and c-MYC with protamine sulfate (5 μg/ml) and incubated overnight. Five days after transduction, the fibroblasts or B-lymphocytes were split onto MEFs and cultured in hiPSC medium for 30 d. hiPSC colonies were manually transferred onto new culture plates and expanded.

### *In vitro* differentiation of neurons, astrocytes and microglia

Glutamatergic neurons were differentiated from hiPSCs as previously described ^27,59^. In brief, hiPSC colonies were detached from the feeder layer using collagenase IV (Thermo) at 37 ℃ for 1 h. 10-16 iPSC colonies (5-8 hiPSC colonies from each well of 24-well plate, 2 wells) were collected in 0.5ml of N2/B27 media, consisting of DMEM/F12, 1× N2 supplement (Gibco), 1× B27 -RA supplement (Gibco) and 1× penicillin–streptomycin, and plated on 35 mm petri dish. On day 1, the medium was changed with N2/B27 medium supplemented with 10 μM SB431542 (Sigma) and 0.1 μM LDN193189 (Stemgent). On day 7, EBs were transferred to Matrigel-coated (10% Matrigel in DMEM/F12, Growth Factor Reduced; Corning) plates and cultured in N2/B27 medium with 10 nM SB431542. On day 16, cells with rosette structures were collected and dissociated mechanically by pipetting. Dissociated cells were plated on Matrigel-coated plates in NPC medium, consisting of DMEM/F12, 1× N2 supplement, 1× B27 -RA supplement, 20 ng/ml basic FGF and 1 μg/ml laminin (Thermo). At 80% confluency, NPC medium was changed into neuronal medium, consisting of DMEM/F12, 1× N2 supplement, 1× B27 -RA supplement, 1× penicillin–streptomycin, 20 ng/ml BDNF (Peprotech), 10 ng/ml GDNF (Peprotech), 250 μg/ml dibutyryl cyclic-AMP (Biogems) and 200 nM L-ascorbic acid (Sigma). The cultures were maintained in neuronal identity was validated by qRT-PCR and immunocytochemistry.

hiPSC-derived astrocytes were generated as previously described ^27,60^. Briefly, hiPSC-derived NPCs were differentiated into astrocytes by seeding dissociated single cells at 15,000 cells/cm^2^ on Matrigel-coated plates in Astrocyte medium (ScienCell: 2% FBS, Astrocyte Growth Supplement, 1% penicillin–streptomycin in astrocyte basal medium) and maintained. After 30– 40 d of differentiation, hiPSC-derived astrocytes were characterized using qRT-PCR and immunocytochemistry.

hiPSC-derived microglia were generated as previously described ^27,61^. Briefly, hiPSC colonies were detached from the feeder layer with collagenase IV for 1 h. hiPSC colonies were collected in microglia medium consisting of 10 ng/ml IL-34 (Peprotech) and 10 ng/ml GM-CSF (Peprotech) in 100-mm sterile petri dishes. After 7–14 d of differentiation, EBs with a cystic morphology were selected and transferred to 10 μg/ml poly-L-ornithine and 5 μg/ml laminin-coated plates. Six additional titrations were performed every 5 d. Further maintenance was performed in microglia medium with 100 ng/ml IL-34 and 5 ng/ml GM-CSF. Microglial identity was validated by qRT-PCR and immunocytochemistry.

### Generation of RELN-expressing neurons

hiPSC-derived RELN-expressing neurons were generated as previously described ^27^. The lentiviral construct for RELN expression was generated by subcloning domains 3–6 (central domains) of *RELN* from the pCrl construct (Addgene #122443, kindly gifted by Mi-Ryoung Song, Gwangju Institute of Science and Technology) into the pLenti 6.3-DEST (Thermo) lentiviral expression vector. NPCs were transduced with the *RELN*-containing lentivirus using protamine sulfate (5 μg/ml). Three days after transduction, NPCs were expanded in NPC medium, containing blasticidin (6 μg/ml), for antibiotic selection. NPCs were then differentiated into glutamatergic neurons. qRT-PCR was performed to confirm RELN expression.

### A stepwise development of forebrain assembloids

Forebrain assembloids were generated as previously described ^27^. hiPSCs were cultured on a feeder layer in a 24-well plate (5-8 hiPSC colonies in each well of the 24-well plate). When the diameter of hiPSC colonies reached 1.0–1.5 mm, the hiPSC medium was replaced with 0.25 ml of 1 mg/ml collagenase IV and incubated at 37 °C for 1-2 h to detach the colonies from the feeder layer without disrupting their colony structure. Detached colonies were transferred to a 15-ml tube and washed with 1 ml of hiPSC medium. 2 ml of EB medium, comprising hiPSC medium (without bFGF) supplemented with 2 μM Dorsomorphin (Sigma) and 2 μM A83-01 (Tocris), was added, and the 5-8 hiPSC colonies (from each well of the 24-well plate) in EB medium were transferred and cultured in a 35-mm petri dish at 37 ℃ to form EBs. On day 5, half of the medium was replaced with neural induction medium consisting of DMEM/F12, supplemented with 1× N2 supplement, 10 μg/ml Heparin (Sigma), 1× penicillin–streptomycin, 1× Non-essential amino acids, 1× Glutamax, 1 μM CHIR99021 (Tocris), and 1 μM SB-431542 (Cellagentech). On day 7, 5-8 EBs were transferred to a 1.5 ml microcentrifuge tube and mixed with 50 μl of Matrigel and 33 μl of neural induction medium. A total of 88 μl of mixture containing EBs was plated onto the center of a 35-mm petri dish and incubated at 37 ℃ for 30 min. Then, 2 ml of neural induction medium was added, and EBs were cultured until day 14 to induce neuroepithelium-like structures. On day 14, the Matrigel-embedded neuroepithelium structures were mechanically dissociated by gentle pipetting, and the resulting structures were then transferred to a new 35-mm petri dish containing 2 ml of differentiation medium consisting of DMEM/F12 supplemented with 1× N2 supplement, 1× B27 supplement, 1× penicillin– streptomycin, 1× 2-Mercaptoethanol, 1× Non-essential amino acids, 1× Glutamax, and 2.5 μg/ml Insulin (Sigma). These neuroepithelium structures were cultured in differentiation medium until day 25 in a shaking incubator (Eppendorf; New Brunswick S41i, 90 rpm) to form forebrain organoids with multiple rosettes.

From days 25 to 32, the organoids were cultured in differentiation medium supplemented with 1 μM CHIR99021 and 400 nM SAG (Millipore). Rosette structures in 32-day-old forebrain organoids were observed under an inverted microscope at 10X magnification (Evos XL Core, Invitrogen), and manually dissected using fine forceps (Fine Science Tools). About 5-7 single-rosette organoids (about 25-40 single-rosette organoids derived from SCZ patients) were generated from one forebrain organoid. Since each batch of hiPSCs typically yields approximately 5-8 forebrain organoids, this results in the generation of approximately 25-56 single-rosette organoids per batch (around 125-320 single-rosette organoids derived from SCZ patients). After dissection, the single-rosette organoids were cultured for an additional 3 days to stabilize the dissected structures.

On day 35, five single-rosette organoids, whose diameters were closest to the median values within the range of 25-40 single-rosette organoids, were selected for further procedures. The resulting single-rosette organoids were then encapsulated with 1–2 μl of Matrigel containing RELN-expressing neurons at a concentration of 1 × 10^4^ cells/μl and cultured in 2 ml of differentiation medium until day 50.

On day 50, hiPSC-derived astrocytes (1 × 10^4^ cells) and microglia (2 × 10^3^ cells) ^62^ were then microinjected into the outer cortical layer of 50-day-old forebrain assembloids. Cells were microinjected at six different locations, which are evenly distributed through the outer cortical layer, with 50 μm inside the surface of the cortical layer. The resulting assembloids were cultured for an additional 30 days in 2 ml of maturation medium consisting of Neurobasal medium (Gibco) supplemented with 1× B27 supplement, 1× penicillin–streptomycin, 1× 2-Mercaptoenthanol, 0.2 mM Ascorbic Acid, 20 ng/ml BDNF (Peprotech), 20 ng/ml GDNF (Peprotech), and 0.5 mM cAMP (Sigma) to generate the final form of forebrain assembloids. From days 14 to 80, all cultures were maintained in a shaking incubator with medium changes every other day.

### Generation of combinatorial, mix-and-match forebrain assembloids

Similar to the overall process of creating normal forebrain assembloids as described above, combinational assembloids were generated by reconstituting organoids derived from healthy individuals (H) or SCZ patients (S) with astrocytes and microglia derived from healthy individuals or SCZ patients. In this way, 8 combinatorial assembloids were created in this study (Organoid-Astrocyte-Microglia; HHH, HHS, HSH, HSS, SHH, SHS, SSH, SSS). For combinatorial assembloids from multiple healthy and SCZ individuals, organoids and glial cells from healthy individual #1 were matched with organoids and glial cells from SCZ #1 to generate mix-and-match assembloid #1. Combinatorial, mix-and-match assembloids #2 - #3 were created in the same way.

### Generation of genetically manipulated hiPSCs

#### Knockout lines

Guide RNAs were designed using CRISPR Design (https://chopchop.cbu.uib.no/) and cloned into pL-CRISPR-EFS-GFP (Addgene #57818). hiPSCs were maintained on Matrigel-coated plates at 80% confluence before electroporation. After electroporation, 30 GFP-positive colonies were picked up and expanded. Gene knockout in each colony was confirmed using qRT-PCR and the Surveyor mutation detection kit and sequencing (IDT Cat. no. 706020). This process was repeated to generate engineered hiPSCs with multiple gene knock-outs. Brain organoids, astrocytes and microglia were differentiated from these genetically manipulated hiPSC lines for further use as indicated. gRNAs sequences used in this study are as follows:

- *TP53*: CCATTGTTCAATATCGTCCG
- *NFATC4*: CTGATGGTCCAAGCCCCGG
- *UCN*: CAGACTCGGGTCCTGGACCC
- *PTPRF*: GGGTTCCCTTCCATCGACAT
- *WNT11*: CATACACGAAGGCCGACTCC
- *THBS4*: GATTGTGAACGGAATCCACC

#### Expression lines

Lentiviral expression vector, pLX313-TP53-WT (118014, Addgene), was used to produce the lentivirus containing TP53 gene. TP53 was replaced with either NFATC4, WNT11, or THBS4 using NheI and SpeI restriction sites to generate lentiviral expression constructs containing these genes. hiPSCs were transduced with resulting lentiviruses and expanded under antibiotic selection. Five colonies were picked and expanded, and gene expression was confirmed by qRT-PCR. Gene-expressing brain organoids were generated from engineered hiPSCs. For the genetic manipulation of astrocytes and microglia, progenitor cells of different lineages were transduced with a lentivirus and expanded under antibiotic selection. Five colonies were picked and expanded, and gene expression was confirmed by qRT-PCR. Manipulated progenitor cells were further differentiated into astrocyte and microglia.

### Lentivirus and retrovirus production

Lentivirus production was performed as previously described ^26,27^. In brief, transfection mixtures were prepared by mixing 9 μg of packaging vector (gag, pol; pCMV.dR 8.74), 3 μg of envelope vector (VSV-G; pMD2.G), 10 μg of the transfer vector of interest and three volumes of TransIT-LT1 Transfection Reagent (Mirus). 48 h after transfection, lentivirus-containing supernatants were collected and filtered through a 0.45-μm filter. The virus-containing supernatants were further concentrated by centrifuging at 24,000 rpm for 2 h at 4 °C.

Retrovirus production was performed as previously described ^26,27^. In brief, transfection mixtures were prepared by mixing 6.5 μg of packaging vector (Addgene #8454), 3.5 μg of envelope vector (Addgene #8449), 10 μg of transfer vector of interest, and three volumes of TransIT-LT1 Transfection Reagent, followed by vortexing with Opti-MEM (Gibco) and the mixture was incubated for 20 min at room temperature (RT). All other procedures were performed as described above for production of lentivirus.

### Electroporation

hiPSCs were treated with the ROCK inhibitor Y27632 (10 μM) 1 h prior to electroporation. Accutase (Sigma)-dissociated single cells were resuspended with Opti-MEM. 10μg of DNA plasmid were delivered to 1 × 10^6^ cells using a NEPA21 electroporator (Poring pulse: 125 V voltage, 2.5 ms pulse length, 50 ms pulse gap, 2 pulses, 10% pulse decay, +orientation; Transfer pulse: 20 V voltage, 50 ms pulse length, 50 ms pulse gap, 5 pulses, 40% pulse decay, ±orientation). Cells were then cultured in mTeSR (StemCell Technologies) containing the ROCK inhibitor Y27632 for the following 48 h.

### qRT-PCR

Total RNA was extracted from multiple cells and organoids as previously described ^26,27^. In brief, cells and organoids were homogenized by trituration and trypsinization. RNA was extracted using the RNeasy Plus Mini Kit (QIAGEN) and first-strand cDNA was synthesised using a High-Capacity cDNA Reverse Transcriptase Kit (Applied Biosystems) with oligo dT. qRT-PCR was performed using SYBR Green Supermix (Applied Biosystems) and a One-step Cycler (Applied Biosystems). Gene expression was normalized to the housekeeping gene GAPDH.

### Immunohistochemistry

Immunohistochemistry was performed as previously described ^26,27^. In brief, brain organoids and assembloids were fixed in 4 % paraformaldehyde (PFA) for 15 min and cryopreserved in 30 % sucrose overnight. Samples were embedded in an OCT compound (Sakura) and frozen at −20 ℃. Then, 8–20-μm-thick sections were generated using a cryostat (Leica). Frozen sections were fixed in 4 % PFA for 20 min at 4 ℃, washed with PBS three times and blocked in 2 % goat serum and PBS containing 0.25 % Triton X-100 (PBS-T) for 1 h at RT. The sections were then incubated with primary antibodies diluted in blocking buffer overnight at 4 ℃. The following primary antibodies were used: Ki67 (1:500, Abcam), p-Vim (1:250, MBL), TUJ1 (1:300, BioLegend), SOX2 (1:300, Abcam; 1:100, Santacruz), CTIP2 (1:300, Abcam), CUX2 (1:300, Abcam), SATB2 (1:300, Abcam), TBR1 (1:300, Abcam), MAP2 (1:300, Abcam), GFAP (1:300, Dako), IBAI (1:60, Santacruz), RELN (1:200, MBL), BRN2 (1:300, Santacruz), PSD95 (1:300, Invitrogen), and NANOG (1:200, Abcam). Sections were washed three times with 0.25 % PBS-T and incubated with secondary antibodies (1:1,000, Life Technologies) diluted in blocking buffer for 1 h at RT. The sections were washed with 0.25 % PBS-T and mounted with Prolong Gold mounting reagent (Invitrogen).

For immunocytochemistry, cells were plated on 10 μg/ml poly-L-ornithine and 5 μg/ml laminin-coated coverslips on a 12-well plate. When reached 80% confluence, cells were washed with PBS and fixed in 4 % PFA for 5 min at RT. Cells were washed with PBS three times and blocked for 40 min at RT. Cells were then incubated with diluted primary antibodies for 1 h at RT and washed three times with PBS-T. Cells were then incubated with secondary antibodies diluted in blocking buffer for 40 min at RT. The cells were washed two times with PBS-T and mounted on glass slides.

### Multi-electrode array (MEA) recording

24-well MEA plates (Axion Biosystems, Atlanta, GA, USA) were coated with 10 μg/ml poly-L-ornithine and 5 μg/ml laminin solution prior to assembloid culture. Assembloids were placed on MEA plates and cultured for 2 weeks to be attached to the electrodes with media change in every 3 days. After two weeks of attachment and stabilization, recordings were captured to measure basic parameters such as weighted mean firing rate using a Maestro MEA system and AxIS Software Spontaneous Neural Configuration (Axion Biosystems). Spike detection was carried out using the AxIS software by setting an adaptive threshold at 5 times the noise’s standard deviation estimated for each channel (electrode). Before recording, the plate was left to stabilize for 10 minutes inside the Maestro device.

The electrodes with at least 5 spikes/min were defined as active electrodes. Raster plot and array-wide spike histogram were obtained using Axion Biosystems’ Neural Metrics Tool.

### Calcium imaging

Calcium imaging of forebrain assembloids was performed as previously described ^27^. In brief, forebrain assembloids were incubated with organoid medium containing 1 μM Fluo4-AM (Invitrogen) for 3 h at 37 ℃. Assembloids were washed once with DPBS and incubated in brain organoid medium. Time-lapse image sequences were acquired for 2 min with 1 sec intervals on a Nikon confocal microscope. ΔF/F traces in the selected cells were calculated and shown in the graph (ΔF/F=(F – F0)/F, where F is the fluorescence at given time point and F0 is the minimum fluorescence of each cell). Spontaneous calcium activities were analyzed with ImageJ software.

### FACS isolation

For FACS of neurons, astrocytes, and microglia of forebrain assembloids, forebrain assembloids were dissociated with Accutase for 1 h and washed with assembloid medium. Cells were resuspended in PBS and incubated for 10 min with FVD510 at 4 °C. Cells were then washed with PBS and incubated for 30 min at 4 °C with the following antibodies resuspended in FACS buffer (DPBS with 1% FBS): anti-CD11b-FITC (BioLegend, #101205), anti-CD49f-PE (BD Bioscience, #555736). Cells were filtered through a strainer cap and run on BD FACSAria^TM^ Fusion. Data were analyzed using FlowJo ver.10.8.1.

For FACS of neurons, astrocytes, and microglia of mouse cerebral cortices, mouse brains were dissociated and single cell suspensions were obtained as previously described ^63^. Cells were resuspended in PBS and incubated for 10 min with FVD510 at 4 °C. Cells were then washed with PBS and incubated for 30 min at 4 °C with the following antibodies resuspended in FACS buffer: anti-CD11b-FITC (BioLegend, #101205), anti-Gfap-AF647 (BD Bioscience, #561470). Cells were filtered through a strainer cap and run on BD FACSAria^TM^ Fusion. Data were analyzed using FlowJo ver.10.8.1.

### Co-expression module analysis for post-mortem human brain tissue

To identify SCZ-associated co-expression modules, two data sources were utilized: 1) 73 spatiotemporally regulated gene co-expression modules in human brain development, identified through weighted correlation network analysis (WGCNA) in the previous study ^33^; and 2) differentially expressed genes (DEGs) between healthy controls and SCZ patients from post-mortem brain tissues, obtained from the previous study ^47^ as a part of the PsychENCODE project^32^.

To identify potential associations of co-expression modules with SCZ, DEGs were ranked based on their log2FC values, and gene set enrichment analysis (GSEA) was performed on the 73 spatiotemporal modules using GSEApy version 0.9.4. Modules that showed significant enrichment in relation to DEGs in SCZ (false discovery rate, FDR < 0.05) were defined as SCZ-associated modules. The normalized enrichment scores (NES) resulting from GSEA are shown in Supplementary Fig. 3c. SCZ-associated modules with positive scaled expression in the early time window were subsequently determined as early developmental modules. Cell-type enrichment analysis was performed on early developmental modules to identify NPC-associated modules. Gene ontology (GO) and biological processes enriched in NPC-associated modules were identified by a hypergeometric test using curated gene set signatures from MSigDB (v7.0). TF enrichment analysis was performed on genes within NPC-associated modules by a hypergeometric test using ‘TFs-target genes’ from the Gene Regulatory Network of PsychENCODE^32^ (Elastic Net regression weight cutoff = 0.1). The enrichment p-value from the hypergeometric test was corrected by the Benjamini–Hochberg procedure. The TF enrichment test on differentially expressed genes (DEGs) between healthy individuals-derived and SCZ patients-derived forebrain organoids was conducted using the ChEA3 database web-server application^64^.

### Prediction analysis to identify the signaling axes for cell–cell interactions

The NicheNet platform^34^ was employed to identify reciprocal ligand-receptor interactions between neurons and glial cells. To identify neuronal factors that might induce transcriptomic changes in glial cells, the sender and receiver were defined as assembloid-derived SCZ neurons and assembloid-derived SCZ astrocytes/microglia, respectively. Conversely, to identify glia cell-derived factors that might induce the transcriptomic changes in neurons, the sender and receiver were defined as assembloid-derived SCZ astrocytes/microglia and assembloid-derived SCZ neurons, respectively. DEGs served as inputs for both sender and receiver in the NicheNet platform.

To obtain DEGs in neurons, astrocytes, and microglia in SCZ assembloids, we performed RNA-seq analysis of assembloid-derived neurons, astrocytes, and microglia from healthy individuals (Healthy # 1, 2, 3) and SCZ patients (SCZ #1, 2, 3). Genes with expression levels below a threshold of 1 [log2(average Transcripts Per Million (TPM) + 1) < 1] were filtered out to obtain genes expressed in specific cell types. DEGs were defined as those meeting the criteria of |log2 Fold Change (FC)| > 1 and an adjusted p-value < 0.05 for each cell type.

To obtain DEGs in neurons, astrocytes, and microglia in post-mortem SCZ brains, scRNA-seq data from SCZ patients, obtained from the National Institute of Mental Health Repository & Genomics Resource, a centralized national biorepository for genetic studies of psychiatric disorders (Synapse ID: syn22362009), was analyzed. Average gene expression within each cluster, annotated as neurons (including both glutamatergic and GABAergic types), astrocytes, and microglia, was calculated. As no genes meet the predefined fold-change filter criteria (|log2FC| > 1) in post-mortem brain data, DEGs were determined based on an adjusted p-value < 0.05 for each cell type.

Four distinct scores—Auroc (Area under receiver operating characteristic), Aupr (Area under precision–Recall), Aupr_correct (Area under precision–Recall corrected), and Pearson correlation coefficient scores—representing the extent to which a specific ligand can regulate the expression of a specific target gene, were calculated for each DEG using the NicheNet platform. The regulatory potential score for individual genes was determined as the multiplied value of these four scores, and genes were ranked based on the regulatory potential scores. The top 50 genes for each cell type, upregulated in SCZ patients (log2FC > 0) and showing the highest regulatory potential, were obtained and considered candidate genes responsible for reciprocal interactions between neurons and glial cells.

### RNA sequencing analysis

#### RNA extraction and RNA-seq library construction

Total RNA was extracted using the RNeasy Mini Kit (Qiagen) following the manufacturer’s instructions. RNA-seq libraries were constructed with 1-5μg of RNA using a TruSeq Stranded mRNA kit (Illumina) following the manufacturer’s instructions. RNA-seq was performed using the Illumina NovaSeq 6000 or HiSeq X Ten platform.

#### RNA-seq data processing and DEG analysis

RNA-seq reads were aligned to the human reference genome GRCh37-hg19 using the STAR (v2.7.2b) ^65^ aligner with ENCODE options. Gene expression levels were quantified with the RSEM (v1.2.31) ^66^ package. For differential expression analysis, DEGUST (degust.erc.monash.edu/.), a web-based tool employing ‘limma’ and ‘edgeR’ packages, was utilized. Count data were filtered based on minimum expression levels and counts per million (CPM) thresholds of 0.5 in at least three samples. Filtered count data were normalized with ‘voom’ function from the ‘limma’ package, preparing the data for linear modelling. A linear model was then fitted to the normalized data using ‘lmFit’. The ‘eBayes’ function was applied to obtain statistics for differential expression.

#### GO and Pearson correlation coefficient analyses

GO enrichment analysis on DEGs was performed using the Geneontology website application (http://geneontology.org/) ^67^. The Pearson correlation coefficients were calculated between the expression matrices of each dataset.

#### Similarity analysis between MAM-treated mice and human SCZ patients

To investigate whether our MAM-treated mouse model may resemble human SCZ patients, RNA-seq on the prefrontal cortex of three MAM-treated mice was performed. We downloaded and compared RNA-seq data from two MAM-treated (GSM2916082 and GSM2916087) and two wild type (GSM2916077 and GSM2916079) mice from a previous study ^37^, along with 338 SCZ and 921 control human samples from PsychEncode project ^68^.

To compare human and mouse RNA expression, 10,519 genes with a Benjamini-Hochberg adjusted p-value of t-test < 0.01 between human SCZ and control samples were selected. Further analysis was performed using 6,726 genes that matched between mouse and official human gene names.

Correlation matrix and an unsupervised cluster was created by calculating the Pearson correlation coefficient between the mouse samples used for analysis. Furthermore, after obtaining the Pearson correlation coefficient between the seven mouse samples and all human samples, the average correlation coefficient between human wild type/SCZ samples and mouse wild type/MAM samples was calculated.

### Mice

8-9 weeks old, male and female C57BL/6 mice were used in all mouse experiments. Mice were housed under specific pathogen-free conditions with regulated daylight (12 h light:dark cycle), humidity, and temperature. All experiments followed all relevant guidelines and regulations. All experiments were approved by the Institutional Animal Care and Use Committee at Seoul National University.

### MAM treatment

MAM (MRIGlobal Chemical Carcinogen Repository, #213) was dissolved in physiological saline. On embryonic day 13 (E13), either saline or MAM (7.5 mg/kg) was administered intraperitoneally (i.p.) to pregnant female mice.

### *In utero* electroporation

pLKO3.G (Addgene, #14748), a gift from Christophe Benoist & Diane Mathis, and pCALSL-miR30 (Addgene, #13786), a gift from Connie Cepko ^69^, were purchased from Addgene. shRNA

sequences targeting genes of interest and scramble sequence that does not match with any sequences of mouse genes were identified from the RNAi Consortium (https://portals.broadinstitute.org/gpp/public/). The shRNA sequence targeting mouse Tp53 and Nfatc4, and scramble sequence were cloned into pLKO.3G. For cell type-specific expressions of shRNA, pCALSL-miR30 and the Cre system were used. pCALSL-miR30, a pri-miRNA-based shRNA-expression vector, contains a lox-stop-lox sequence upstream of miR30, which can be induced by Cre-mediated DNA recombination. The sequence targeting mouse Ucn, Ptprf, Wnt11, Thbs4, and scramble sequence were cloned into pCALSL-miR30. pCAG-Cre-IRES-EGFP was used for promoter-dependent Cre and EGFP expression. CAG promoter was replaced with either hSyn1, hGFAP, or mCd11b promoter for shRNA expression in neurons, astrocytes, and microglia, respectively. shRNA sequences and scramble sequence used in this study are as follows:

- *Tp53*: ACCGCCGTACAGAAGAAGAAA
- *Nfatc4*: GGACGGCTCTCCTAGAGATTA
- *Ucn*: GAGCAGAACCGCATCATATTC
- *Ptprf*: CGCTTTGAGGTAATTGAGTTT
- *Wnt11*: CCCTGGAAACGAAGTGTAAAT
- *Thbs4*: GCAGTAGCAGAACCTGGTATT
- *Scramble*: CCTAAGGTTAAGTCGCCCTCG

To observe the effects Tp53, Nfatc4, Ucn, Ptprf, Wnt11, and Thbs4 knockdown has in brain structure and function, in utero electroporation was performed as described previously ^70,71^, with minor modifications. Briefly, E14 or E17.5 pregnant female mice were anesthetized using isoflurane with a standard vaporizer. The hair in the abdominal region was shaved, and the abdomen was then washed with 70% ethanol and treated with iodine solution. The uterus was carefully taken out following a 2 cm laparotomy in the low middle abdomen. shRNA-containing vectors of interest (1-2μg/μl) mixed with 0.3 % Fast Green were injected (1-3μl) into both lateral ventricles of the embryos. Electric pulses (Poring pulse: voltage, 40 V; pulse length, 30 ms; pulse interval, 450 ms; number of pulses, 3; decay rate, 10 %; polarity, + / Transfer pulse: voltage, 25 V; pulse length, 50 ms; pulse interval, 450 ms; number of pulses, 5; decay rate, 40 %; polarity,

+) were transversely applied to the brains with an electroporator (NEPA21, CUY650P5), on each perpendicular side. For E14 mice, the vectors that express the shRNA sequences targeting Tp53 and Nfatc4 were inserted into the lateral ventricles of the embryos, and electroporation was conducted. In the case of E17.5 mice, the vectors were separately injected depending on the cell type: vectors that express shRNA sequences that target neuronal genes (Ucn and Ptprf) were inserted first, followed by electroporation, and then the vectors that express shRNA sequences that target glial cell genes (Wnt11 and Thbs4) were inserted next, which then were also followed by electroporation. The uterus was then placed back into the abdominal cavity, and the abdomen wall and skin were sutured. The embryos were allowed to develop until they were evaluated.

### Behavior tests

Prior to behavior tests, all mice were handled for at least four days to reduce the stress induced by the experimenters.

#### Sucrose preference test

Sucrose preference test was performed as previously reported ^72^. Briefly, mice were housed individually for two days to acclimate them to drinking water from two bottles. On the day of the test, one of the bottles was filled with 1% sucrose and the other with drinking water, and both bottles were weighed. Twelve hours later, the bottles were rearranged to prevent a side preference. At 24 h, the bottles were weighed and sucrose preference was determined as a percentage of sucrose solution consumed (amount of sucrose solution consumed/total amount of solution consumed x 100).

#### Open field test

Open field test was performed as previously reported ^72^, with minor modifications. Mice were individually placed in a white plastic open-field apparatus (50 cm × 50 cm × 50 cm), where the center zone was defined as a square at the center (25 × 25 cm), illuminated by a white light 2 m overhead (6-8 lux). During the test, mice were consistently placed at the corner of the apparatus from the same direction. The locomotor activity (total distance traveled (cm), time spent in the center area (s), frequency entering the center area) of each animal was recorded for 10 min and analyzed using EthoVision XT.

#### Elevated plus maze test

Elevated plus maze test was performed as previously described ^73^, with minor modifications. Mice were placed in a plus-shaped plastic apparatus, consisting of two open arms (30 cm × 5 cm) and two closed arms (30 cm × 5 cm) with 15-cm-high walls and a central square (5 cm × 5 cm), illuminated by a white light 2 m overhead (6-8 lux). The floor of the arms and central square were elevated to a height of 50 cm above the floor. During the test, mice were consistently placed in one of two closed arms from the same direction. The locomotor activity (total distance traveled (cm), time spent in the open arms (s), frequency entering the open arms) of each animal was recorded for 10 min and analyzed using EthoVision XT.

## Data analysis

Statistical analyses were performed using GraphPad Prism ver. 10. All data were presented as mean +/− SEM. Comparisons between groups were performed using an unpaired *t*-test or a nested *t*-test as described in figure legends. Values of *p* < 0.05 were considered statistically significant.

## Data availability

All data either generated or analyzed during this study are included in this article (and its supplementary information files).

## Code availability

The codes used for data analysis are available upon request.

## Acknowledgements

We thank Mi-Ryoung Song at GIST for their generous provision of the RELN-expressing construct. This research was supported by grants from the National Research Foundation of Korea (NRF-2022R1A2C3002702, RS-2023-00223277), Samsung Science and Technology Foundation (SSTF-BA2101-12), New Faculty Startup Fund from Seoul National University, and the BK21FOUR Research Fellowship.

## Author contributions

E.K, Y.K and K.S conceived the ideas and experimental design. E.K, Y.K, S.H, I.K, and J.L performed the overall experiments. Y.K analyzed RNA-seq data. K.L performed the bioinformatical analyse and S.K provided advice on bioinformatic analyses. M.A and S.K helped with behaviour analyses of SCZ mice. E.K, Y.K and K.S wrote the manuscript.

## Supplementary Figure legends

**Supplementary Fig. 1.**
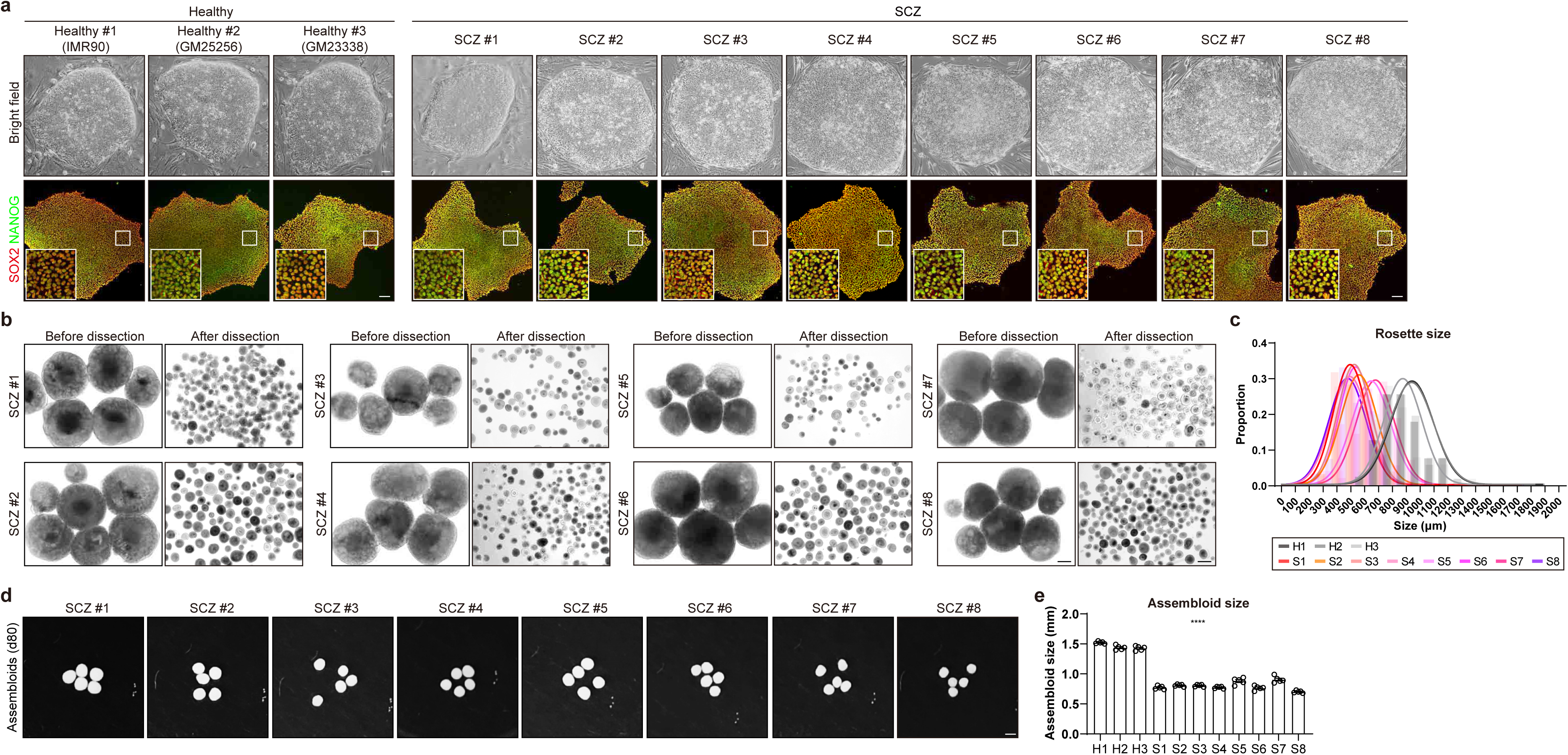
Generation of patient-derived forebrain assembloids from human SCZ patients. (a) Representative images of hiPSC colonies from three healthy individuals and eight SCZ patients. The upper panels show bright field images. The lower panels show immunostaining images for SOX2 and NANOG. Scale bars, 100 μm. (b) Representative images of manually-dissected single rosette organoids from healthy individuals and SCZ patients (left, before dissection; right, after dissection). Scale bars, 1 mm. (c) Quantification of the size of manually-dissected, single rosette organoids. Single rosette organoids were categorized by size based on the length of their diameters. The bar graphs show the relative proportion of the different-sized single rosette organoids. Normal distribution is represented by the curve. (d) Representative images of forebrain assembloids (d80) derived from SCZ patients. Scale bars, 1 mm. (e) Quantification of the size of forebrain assembloids derived from healthy individuals and SCZ patients. The size of the assembloids was determined by averaging the diameters of the assembloids, measured twice at a 90-degree angle (Five biological replicates were evaluated; n = 5). Significance was calculated using a nested *t*-test.

**Supplementary Fig. 2.**
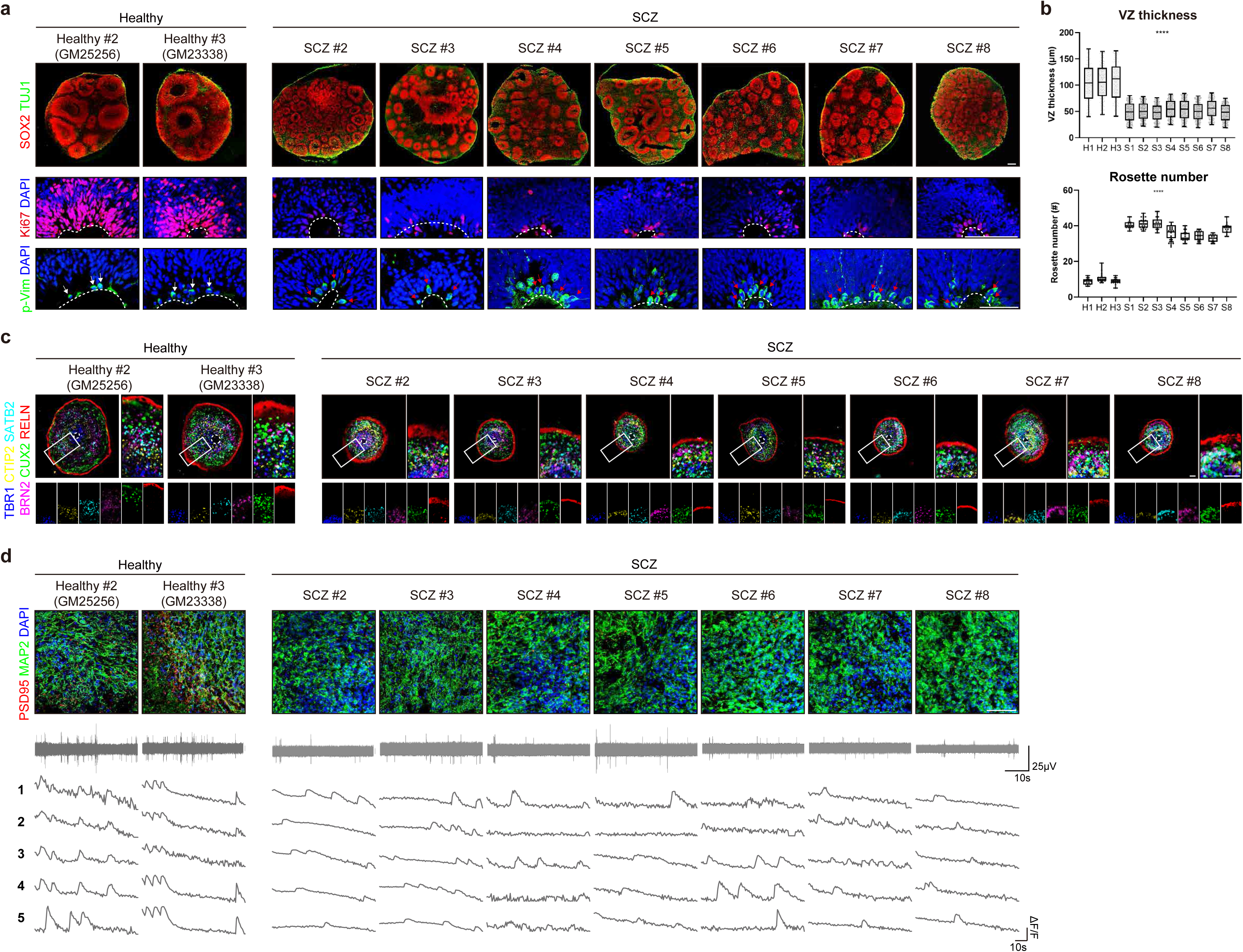
Premature neurogenesis and weakened cortex organization with reduced synaptic connectivity in SCZ patient-derived forebrain assembloids. (a) (top) Representative images of healthy and SCZ brain organoids (d32) immunostained with SOX2 and TUJ1. Scale bars, 100 μm. (middle) Representative images of healthy and SCZ organoids (d32) immunostained with Ki67. Scale bars, 100 μm. (bottom) Representative images of healthy and SCZ organoids (d32) immunostained with p-Vim. The white arrows indicate NPCs that underwent horizontal division; the red arrows indicate NPCs that underwent oblique or vertical division. Scale bars, 50 μm. (b) (top) Quantification of VZ thickness by measuring the thickness of VZ from two independent regions in each rosette (at 90-degree angle intervals). The VZ thickness of all rosettes was quantified in two sections of one organoid. All organoids in each group are counted. n = 156 (H1), n = 272 (H2), n = 161 (H3), n = 820 (S1), n = 874 (S2), n = 802 (S3), n = 736 (S4), n = 744 (S5), n = 766 (S6), n = 648 (S7), n =794 (S8). Significance was calculated using a nested *t*-test. Center line, median; whiskers, min to max (show all points). (bottom) Quantification of the number of rosettes. The number of rosettes was quantified in three sections of one organoid. All organoids in each group are counted. n = 15 (H1), n = 18 (H2), n = 15 (H3), n = 15 (S1), n = 21 (S2), n = 18 (S3), n = 18 (S4), n = 15 (S5), n = 18 (S6), n = 15 (S7), n =15 (S8). Significance was calculated using a nested *t*-test. Center line, median; whiskers, min to max (show all points). (c) Representative images of 6-layered cortical structures of healthy and SCZ assembloids (d80). Upper left panels show merged images of three serial sections at 8-μm intervals in which each section was immunostained for TBR1/CTIP2, SATB2/RELN, and BRN2/CUX2, respectively. Magnified images (inset in the left panels) are shown on the right. Scale bars, 100 μm. (d) (top) Representative images of healthy and SCZ assembloids (d80) immunostained for neurons (MAP2) and synapses (PSD95). Scale bars, 50 μm. (middle) Representative images of a recording plot from a single electrode of a MEA in healthy and SCZ assembloids (d80). (bottom) Representative images of calcium imaging analyses of selected cells in healthy and SCZ assembloids (d80).

**Supplementary Fig. 3.**
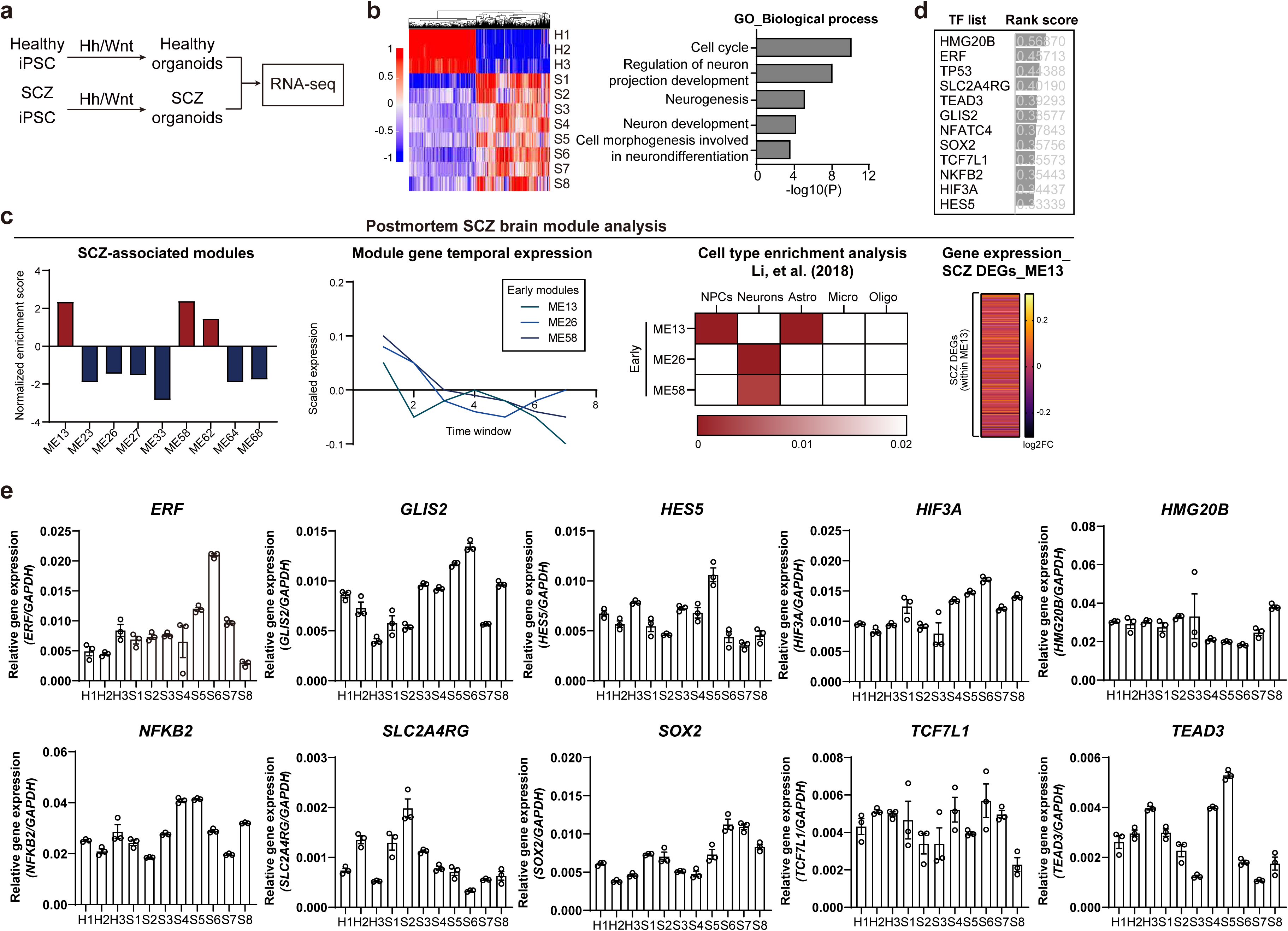
Comparative transcriptome analysis of post-mortem SCZ brains and SCZ-derived forebrain assembloids to identify pathogenic regulators at the early stage of human SCZ brain development. (a) Experimental strategy for RNA-seq analysis of early forebrain organoids (d32) derived from healthy individuals and SCZ patients. (b) (left) Heatmap representation of RNA-seq analysis of forebrain organoids derived from healthy individuals and SCZ patients. (right) GO analysis of DEGs between forebrain organoids derived from healthy individuals and SCZ patients using selected DEGs (FDR < 0.01) is shown on the right. (c) (left) Normalized enrichment scores of nine schizophrenia (SCZ)-associated modules (ME13, ME58 and ME62; upregulated and ME23, ME26, ME27, ME33, ME64 and ME68; downregulated). (middle left) Identification of three early modules (ME13, ME26 and ME58) by analyzing temporal gene expression in each module. (middle right) Cell-type enrichment analysis of three early modules. (right) Log2FC value of SCZ DEGs within NPC module (ME13). Among the 490 genes composing the ME13 module, log2FC values for 480 genes corresponding to SCZ DEGs identified from post-mortem brain analysis are presented. (d) The expression of twelve candidate TFs, which were identified in the co-expression network analysis of post-mortem brain tissues, in DEGs between forebrain organoids derived from healthy individuals and SCZ patients are shown along with their rank scores. The rank score is calculated as 1/log(score), where the score obtained from a TF enrichment analysis conducted on the ChEA3 platform. (e) Relative gene expression of *ERF, GLIS2, HES5, HIF3A, HMG20B, NFKB2, SLC2A4RG, SOX2*, *TCF7L1,* and *TEAD3* in early forebrain organoids derived from healthy individuals and SCZ patients (Three biological replicates were evaluated; n = 3). Significance was calculated using a nested *t*-test.

**Supplementary Fig. 4.**
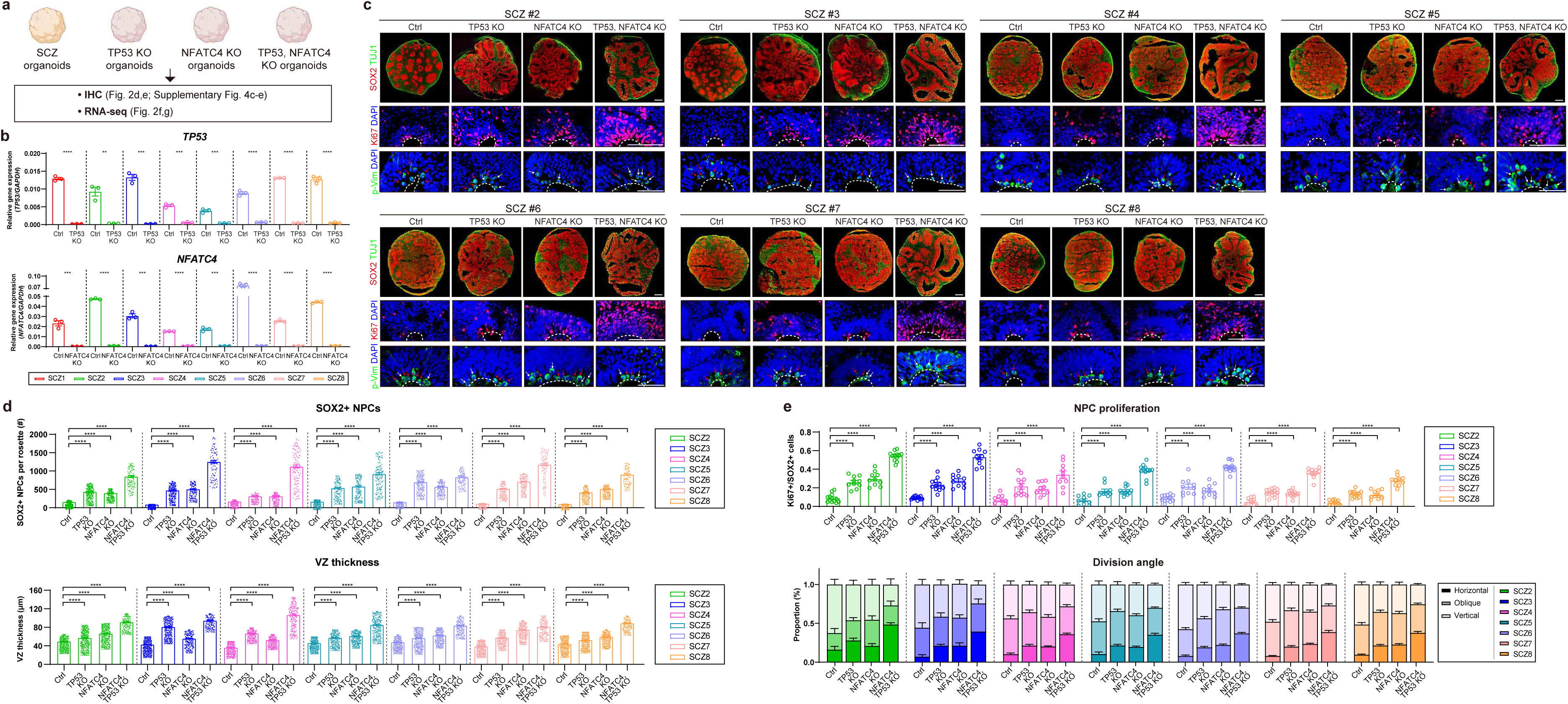
Alterations in the transcriptome landscape with enhanced proliferation and decreased differentiation of NPCs in TP53/NFATC4-ablated organoids from schizophrenia patients. (a) Diagram of the experimental design to investigate the role of TP53 and NFATC4 by generating TP53- and NFATC4-knockout (KO) organoids (d32) derived from SCZ patients. (b) Relative gene expressions of *TP53* and *NFATC4* in TP53- and NFATC4-knockout forebrain organoids from SCZ patients (Three biological replicates were evaluated; n = 3). Significance was calculated using an unpaired *t*-test. (c) (top) Representative images of genetically manipulated SCZ forebrain organoids immunostained with SOX2 and TUJ1. Scale bars, 100 μm. (middle) Representative images of genetically manipulated SCZ forebrain organoids immunostained with Ki67. Scale bars, 100 μm. (bottom) Representative images of genetically manipulated SCZ forebrain organoids immunostained with p-Vim. The white arrows indicate NPCs that underwent horizontal division; the red arrows indicate NPCs that underwent oblique or vertical division. Scale bars, 50 μm. (d) (top) Quantification of the NPC population by counting the number of SOX2-positive cells in each rosette. The number of NPCs in all rosettes was quantified in two sections of one organoid. All organoids in each group are counted. S2-derived organoids: n = 380 (Ctrl), n = 299 (TP53-KO), n = 240 (NFATC4-KO), n = 110 (TP53/NFATC4-KO); S3-derived organoids: n = 479 (Ctrl), n = 296 (TP53-KO), n = 169 (NFATC4-KO), n = 98 (TP53/NFATC4-KO); S4-derived organoids: n = 379 (Ctrl), n = 215 (TP53-KO), n = 248 (NFATC4-KO), n = 131 (TP53/NFATC4-KO); S5-derived organoids: n = 458 (Ctrl), n = 183 (TP53-KO), n = 214 (NFATC4-KO), n = 170 (TP53/NFATC4-KO); S6-derived organoids: n = 374 (Ctrl), n = 277 (TP53-KO), n = 256 (NFATC4-KO), n = 139 (TP53/NFATC4-KO); S7-derived organoids: n = 480 (Ctrl), n = 226 (TP53-KO), n = 260 (NFATC4-KO), n = 125 (TP53/NFATC4-KO); S8-derived organoids: n = 348 (Ctrl), n = 286 (TP53-KO), n = 254 (NFATC4-KO), n = 118 (TP53/NFATC4-KO). Significance was calculated using an unpaired *t*-test. (bottom) Quantification of VZ thickness by measuring the thickness of VZ from two independent regions in each rosette (at 90-degree angle intervals). The VZ thickness of all rosettes was quantified in two sections of one organoid. All organoids in each group are counted. S2-derived organoids: n = 760 (Ctrl), n = 598 (TP53-KO), n = 480 (NFATC4-KO), n = 220 (TP53/NFATC4-KO); S3-derived organoids: n = 958 (Ctrl), n = 592 (TP53-KO), n = 338 (NFATC4-KO), n = 196 (TP53/NFATC4-KO); S4-derived organoids: n = 758 (Ctrl), n = 430 (TP53-KO), n = 496 (NFATC4-KO), n = 262 (TP53/NFATC4-KO); S5-derived organoids: n = 916 (Ctrl), n =366 (TP53-KO), n = 428 (NFATC4-KO), n = 340 (TP53/NFATC4-KO); S6-derived organoids: n = 748 (Ctrl), n = 554 (TP53-KO), n = 512 (NFATC4-KO), n = 278 (TP53/NFATC4-KO); S7-derived organoids: n = 960 (Ctrl), n = 452 (TP53-KO), n = 520 (NFATC4-KO), n = 250 (TP53/NFATC4-KO); S8-derived organoids: n = 696 (Ctrl), n = 572 (TP53-KO), n = 508 (NFATC4-KO), n = 236 (TP53/NFATC4-KO). Significance was calculated using an unpaired *t*-test. (e) (top) Quantification of NPC proliferation by counting Ki67-positive cells as a percentage of the total SOX2-positive cells in each section. NPC proliferation was quantified in two sections of one organoid. All organoids in each group are counted. S2-derived organoids: n = 12 (Ctrl), n = 10 (TP53-KO), n = 10 (NFATC4-KO), n = 12 (TP53/NFATC4-KO); S3-derived organoids: n = 14 (Ctrl), n = 12 (TP53-KO), n = 10 (NFATC4-KO), n = 10 (TP53/NFATC4-KO); S4-derived organoids: n = 10 (Ctrl), n = 12 (TP53-KO), n = 10 (NFATC4-KO), n = 10 (TP53/NFATC4-KO); S5-derived organoids: n = 10 (Ctrl), n =10 (TP53-KO), n = 12 (NFATC4-KO), n = 12 (TP53/NFATC4-KO); S6-derived organoids: n = 12 (Ctrl), n = 10 (TP53-KO), n = 10 (NFATC4-KO), n = 14 (TP53/NFATC4-KO); S7-derived organoids: n = 10 (Ctrl), n = 14 (TP53-KO), n = 12 (NFATC4-KO), n = 10 (TP53/NFATC4-KO); S8-derived organoids: n = 12 (Ctrl), n = 12 (TP53-KO), n = 10 (NFATC4-KO), n = 10 (TP53/NFATC4-KO). Significance was calculated using an unpaired *t*-test. (bottom) The proportion of NPCs with the horizontal, oblique, or vertical orientation of cell division in each section (0–30°: horizontal, 30–60°: oblique, and 60– 90°: vertical). The proportion of horizontal, oblique, or vertical division was quantified based on the average division angle of cells in each section. Two sections were quantified for one organoid. All organoids in each group are counted. S2-derived organoids: n = 12 (Ctrl), n = 10 (TP53-KO), n = 10 (NFATC4-KO), n = 12 (TP53/NFATC4-KO); S3-derived organoids: n = 14 (Ctrl), n = 12 (TP53-KO), n = 10 (NFATC4-KO), n = 10 (TP53/NFATC4-KO); S4-derived organoids: n = 10 (Ctrl), n = 12 (TP53-KO), n = 10 (NFATC4-KO), n = 10 (TP53/NFATC4-KO); S5-derived organoids: n = 10 (Ctrl), n =10 (TP53-KO), n = 12 (NFATC4-KO), n = 12 (TP53/NFATC4-KO); S6-derived organoids: n = 12 (Ctrl), n = 10 (TP53-KO), n = 10 (NFATC4-KO), n = 14 (TP53/NFATC4-KO); S7-derived organoids: n = 10 (Ctrl), n = 14 (TP53-KO), n = 12 (NFATC4-KO), n = 10 (TP53/NFATC4-KO); S8-derived organoids: n = 12 (Ctrl), n = 12 (TP53-KO), n = 10 (NFATC4-KO), n = 10 (TP53/NFATC4-KO).

**Supplementary Fig. 5.**
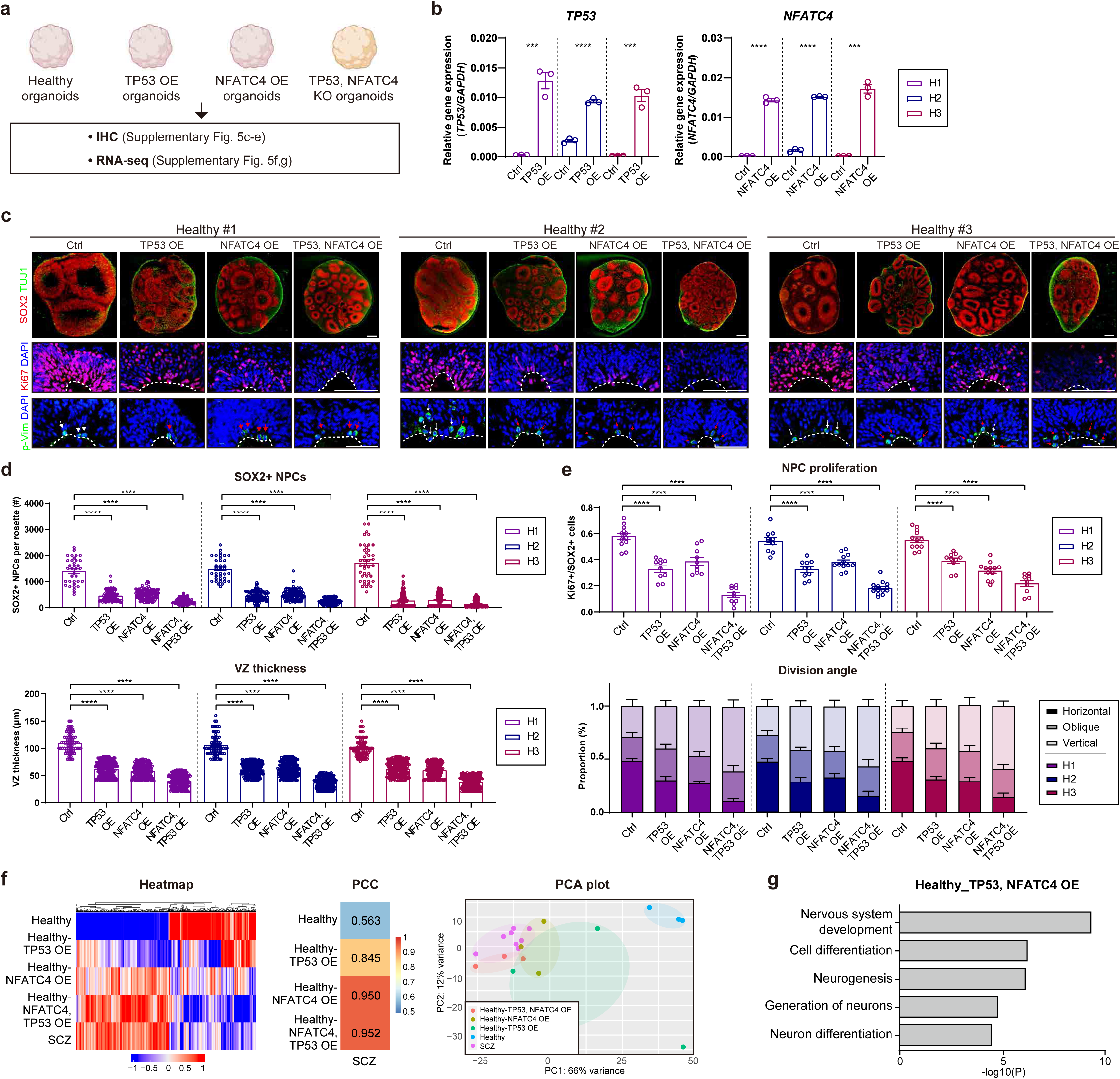
Alterations in the transcriptome landscape with reduced proliferation and increased differentiation of NPCs in TP53/NFATC4-overexpressing organoids from healthy individuals. (a) Diagram of the experimental design to investigate the role of TP53 and NFATC4 by generating TP53- and NFATC4-overexpressing (OE) organoids (d32) derived from healthy individuals. (b) Relative gene expression of *TP53* and *NFATC4* in TP53- and NFATC4-overexpressed healthy forebrain organoids (Three biological replicates were evaluated; n = 3). Significance was calculated using an unpaired *t*-test. (c) (top) Representative images of genetically manipulated healthy forebrain organoids immunostained with SOX2 and TUJ1. Scale bars, 100 μm. (middle) Representative images of genetically manipulated healthy forebrain organoids immunostained with Ki67. The dotted lines demarcate the border between the VZ and the ventricle. Scale bars, 100 μm. (bottom) Representative images of genetically manipulated healthy forebrain organoids immunostained with p-Vim. The dotted lines demarcate the border between the VZ and the ventricle. The white arrows indicate NPCs that underwent horizontal division; the red arrows indicate NPCs that underwent oblique or vertical division. Scale bars, 50 μm. (d) (top) Quantification of the NPC population by counting the number of SOX2-positive cells in each rosette. The number of NPCs in all rosettes was quantified in two sections of one organoid. All organoids in each group are counted. H1-derived organoids: n = 35 (Ctrl), n = 117 (TP53-KO), n = 108 (NFATC4-KO), n = 133 (TP53/NFATC4-KO); H2-derived organoids: n = 36 (Ctrl), n = 107 (TP53-KO), n = 105 (NFATC4-KO), n = 144 (TP53/NFATC4-KO); H3-derived organoids: n = 41 (Ctrl), n = 103 (TP53-KO), n = 106 (NFATC4-KO), n = 136 (TP53/NFATC4-KO). Significance was calculated using an unpaired *t*-test. (bottom) Quantification of VZ thickness by measuring the thickness of VZ from two independent regions in each rosette (at 90-degree angle intervals). The VZ thickness of all rosettes was quantified in two sections of one organoid. All organoids in each group are counted. H1-derived organoids: n = 70 (Ctrl), n = 234 (TP53-KO), n = 216 (NFATC4-KO), n = 266 (TP53/NFATC4-KO); H2-derived organoids: n = 72 (Ctrl), n = 214 (TP53-KO), n =210 (NFATC4-KO), n = 288 (TP53/NFATC4-KO); H3-derived organoids: n = 82 (Ctrl), n = 206 (TP53-KO), n = 212 (NFATC4-KO), n = 272 (TP53/NFATC4-KO). Significance was calculated using an unpaired *t*-test. (e) (top) Quantification of NPC proliferation by counting Ki67-positive cells as a percentage of the total SOX2-positive cells in each section. NPC proliferation was quantified in two sections of one organoid. All organoids in each group are counted. H1-derived organoids: n = 12 (Ctrl), n = 10 (TP53-KO), n = 10 (NFATC4-KO), n = 10 (TP53/NFATC4-KO); H2-derived organoids: n = 10 (Ctrl), n = 10 (TP53-KO), n =12 (NFATC4-KO), n = 14 (TP53/NFATC4-KO); H3-derived organoids: n = 12 (Ctrl), n = 10 (TP53-KO), n = 12 (NFATC4-KO), n = 10 (TP53/NFATC4-KO). Significance was calculated using an unpaired *t*-test. (bottom) The proportion of NPCs with the horizontal, oblique, or vertical orientation of cell division in each section (0–30°: horizontal, 30–60°: oblique, and 60–90°: vertical). The proportion of horizontal, oblique, or vertical division was quantified based on the average division angle of cells in each section. Two sections were quantified for one organoid. All organoids in each group are counted. H1-derived organoids: n = 12 (Ctrl), n = 10 (TP53-KO), n = 10 (NFATC4-KO), n = 10 (TP53/NFATC4-KO); H2-derived organoids: n = 10 (Ctrl), n = 10 (TP53-KO), n =12 (NFATC4-KO), n = 14 (TP53/NFATC4-KO); H3-derived organoids: n = 12 (Ctrl), n = 10 (TP53-KO), n = 12 (NFATC4-KO), n = 10 (TP53/NFATC4-KO). (n) RNA-seq analysis of control, TP53-OE, NFATC4-OE, and both TP53- and NFATC4-OE organoids derived from healthy individuals, as well as organoids derived from SCZ patients. (f) (left) Heatmap representation of RNA-seq analysis of indicated forebrain organoids using selected DEGs (log2 fold change >1, p-value < 0.001). (middle) Pearson correlation coefficient obtained by using selected DEGs (log2 fold change >1, p-value < 0.001) of each forebrain organoid. (right) PCA plot from RNA-seq analysis of indicated forebrain organoids analyzed by using selected DEGs (log2 fold change >1, p-value < 0.001). (g) GO analysis of TP53- and NFATC4-overexpressed forebrain organoids using selected DEGs (FDR < 0.01).

**Supplementary Fig. 6.**
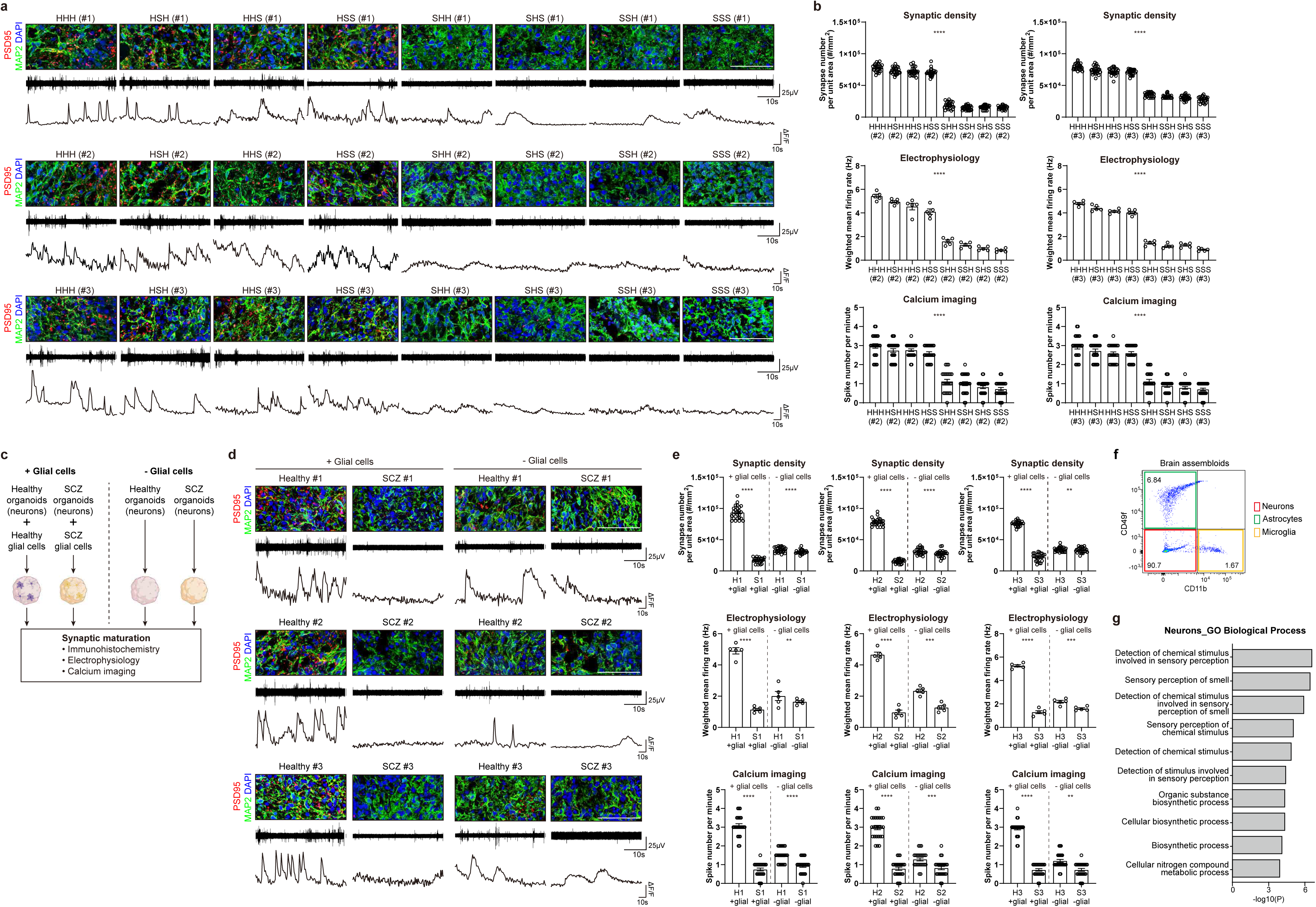
Functional analyses of combinational, mix-and-match forebrain assembloids to show the neuron-dependent, transcriptional plasticity of glial cells in human SCZ brain. (a) (top) Immunostaining analysis for synaptic density of eight combinational assembloids derived from healthy individuals and SCZ patients (Healthy #1 and SCZ #1, Healthy #2 and SCZ #2, and Healthy #3 and SCZ #3). Scale bars, 50 μm. (middle) Representative images of a recording plot from a single electrode of a MEA in eight combinational assembloids derived from healthy individuals and SCZ patients (Healthy #1 and SCZ #1, Healthy #2 and SCZ #2, and Healthy #3 and SCZ #3). (bottom) Representative images of calcium imaging analysis of selected cells in eight combinational assembloids derived from healthy individuals and SCZ patients (Healthy #1 and SCZ #1, Healthy #2 and SCZ #2, and Healthy #3 and SCZ #3). (b) (top) Quantification of the synaptic density in the indicated combinatorial assembloids derived from healthy individuals and SCZ patients (left, Healthy #2 and SCZ #2; right, Healthy #3 and SCZ #3) by calculating the number of synapses per unit area (mm^2^) in each section. Five sections were quantified for one assembloid (Five biological replicates were evaluated; n = 25). Significance was calculated using a nested *t*-test. (middle) Quantification of the weighted mean firing rate in the indicated combinatorial assembloids derived from healthy individuals and SCZ patients (left, Healthy #2 and SCZ #2; right, Healthy #3 and SCZ #3) measured by MEA. Recordings were performed for thirty minutes. The signals from all active electrodes were quantified for each assembloid (Five biological replicates were evaluated; n = 5). Significance was calculated using a nested *t*-test. (bottom) Quantification of spike number per minute analyzed by calcium imaging of selected cells in indicated combinatorial assembloids derived from healthy individuals and SCZ patients (left, Healthy #2 and SCZ #2; right, Healthy #3 and SCZ #3). The number of spikes was quantified in five selected cells of one assembloid (Five biological replicates were evaluated; n = 25). Significance was calculated using a nested *t*-test. (c) Experimental scheme for functional analyses of forebrain assembloids in the presence or absence of glial cells. (d) (top) Immunostaining analysis for synaptic density of healthy and SCZ forebrain assembloids with or without glial cells. Scale bars, 50 μm. (middle) Representative images of a recording plot from a single electrode of a MEA in healthy and SCZ forebrain assembloids with or without glial cells. (bottom) Representative images of calcium imaging analysis of selected cells in healthy and SCZ forebrain assembloids with or without glial cells. (e) (top) Quantification of the synaptic density in healthy and SCZ forebrain assembloids with or without glial cells by calculating the number of synapses per unit area (mm^2^) in each section. Five sections were quantified for one assembloid (Five biological replicates were evaluated; n = 25). Significance was calculated using an unpaired *t*-test. (middle) Quantification of the weighted mean firing rate in healthy and SCZ forebrain assembloids with or without glial cells measured by MEA. Recordings were performed for thirty minutes. The signals from all active electrodes were quantified for each assembloid (Five biological replicates were evaluated; n = 5). Significance was calculated using an unpaired *t*-test. (bottom) Quantification of spike number per minute analyzed by calcium imaging of selected cells in healthy and SCZ forebrain assembloids with or without glial cells. The number of spikes was quantified in five selected cells of one assembloid (Five biological replicates were evaluated; n = 25). Significance was calculated using an unpaired *t*-test. (f) FACS plots gated on neurons (CD49f-CD11b-), astrocytes (CD49f+ CD11b-), and microglia (CD49f-CD11b+) isolated from forebrain assembloids. (g) GO analysis for DEGs between neurons isolated from healthy and SCZ forebrain assembloids using selected DEGs (FDR < 0.01).

**Supplementary Fig. 7.**
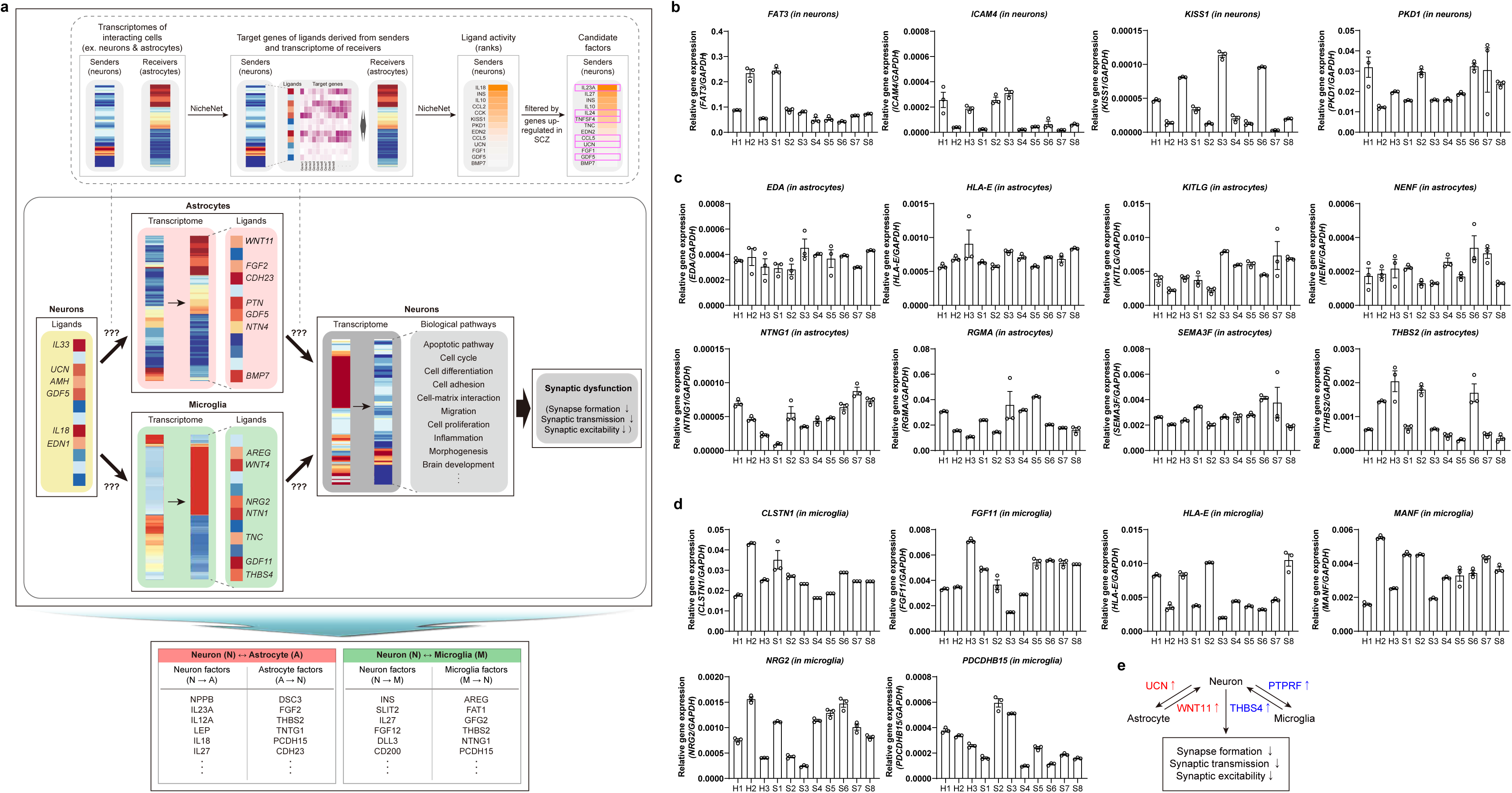
Identification of the UCN/PTPRF-WNT11/THBS4 signaling axis between neurons and glial cells at the late stage of SCZ brain development. (a) Experimental strategy of identifying common factors involved in reciprocal signaling between neurons and glial cells in schizophrenic forebrain assembloids and post-mortem brains. (b-d) Relative expressions of genes in neurons (*FAT3, ICAM4, KISS1,* and *PKD1*) (b), astrocytes (*EDA, HLA-E, KITLG, NENF, NTNG1, RGMA, SEMA3F,* and *THBS2*) (c), and microglia (*CLSTN1, FGF11, HLA-E, MANF, NRG2,* and *PCDHB15*) (d), isolated from healthy and schizophrenic forebrain assembloids (Three biological replicates were evaluated; n = 3). Significance was calculated using a nested *t*-test. (e) Schematic illustration of the proposed model of the dynamic interplay between neurons and glial cells.

**Supplementary Fig. 8.**
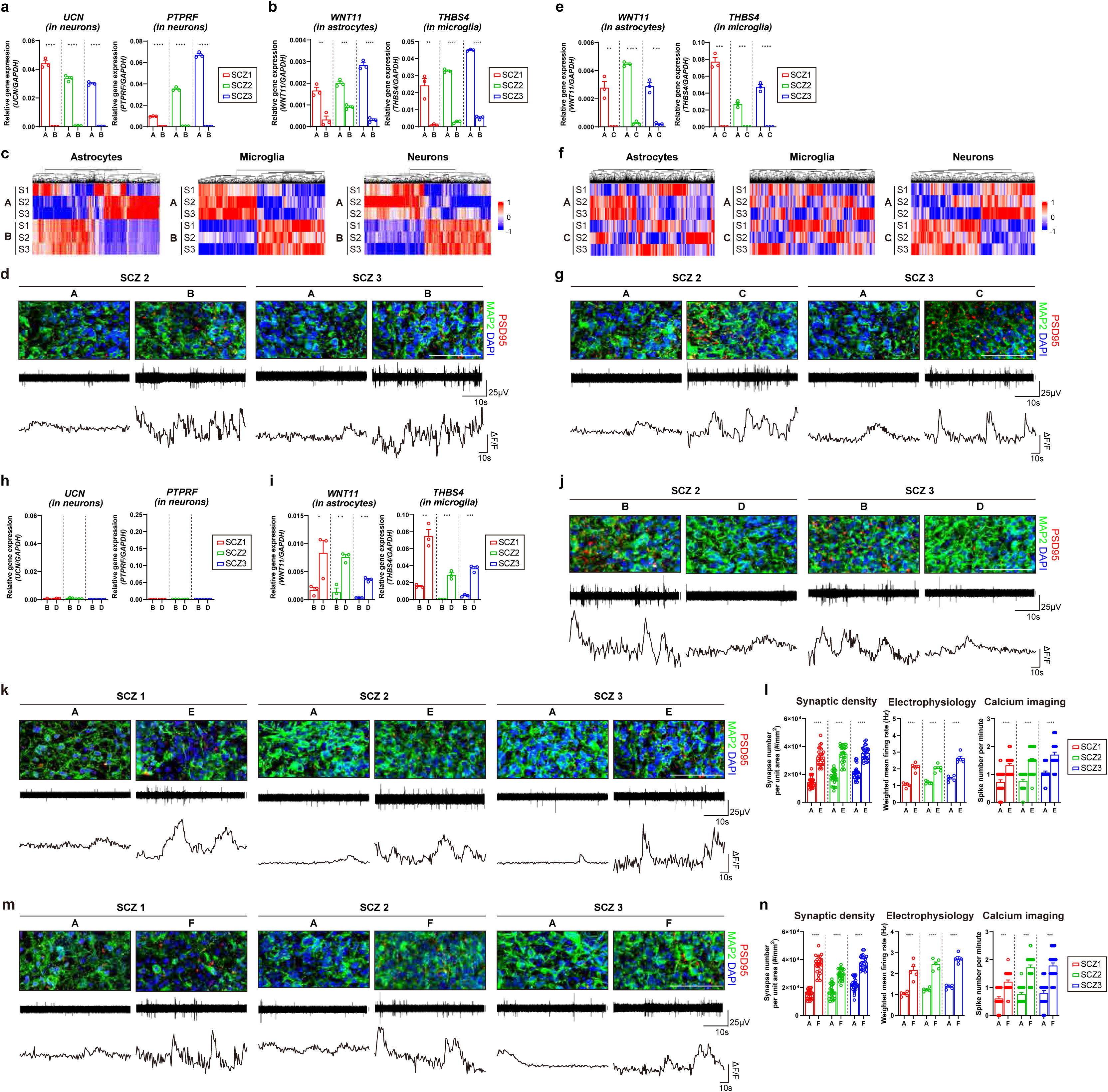
Late developmental defects in the SCZ brain caused by a heightened activity of the UCN/PTPRF-WNT11/THBS4 signaling axis between neurons and glial cells. (a, h) Relative expressions of *UCN* and *PTPRF* in neurons isolated from genetically manipulated forebrain assembloids as indicated in Fig. 4a (a) or Fig. 4g (h). Three biological replicates were evaluated (n = 3). Significance was calculated using an unpaired *t*-test. (b, e, i) Relative expressions of *WNT11* and *THBS4* in astrocytes and microglia, respectively, isolated from genetically manipulated forebrain assembloids as indicated in Fig. 4a (b), Fig. 4d (e), or Fig. 4g (i). Three biological replicates were evaluated (n = 3). Significance was calculated using an unpaired *t*-test. (c, f) Heatmap representation of RNA-seq analysis for neurons, astrocytes, and microglia isolated from genetically manipulated forebrain assembloids as indicated in Fig. 4a (c) or Fig. 4d (f). (d, g, j) (top) Immunostaining analysis of synaptic density of indicated forebrain assembloids derived from SCZ patients (SCZ #2 and #3). Scale bars, 50 μm. (middle) Representative images of a recording plot from a single electrode of a MEA in indicated forebrain assembloids derived from SCZ patients (SCZ #2 and #3). (bottom) Representative images of calcium imaging analysis of selected cells in indicated forebrain assembloids derived from SCZ patients (SCZ #2 and #3). (k, m) (top) Immunostaining analysis of synaptic density of indicated forebrain assembloids derived from SCZ patients (SCZ #1, 2, 3). Scale bars, 50 μm. (middle) Representative images of a recording plot from a single electrode of a MEA in indicated forebrain assembloids derived from SCZ patients (SCZ #1, 2, 3). (bottom) Representative images of calcium imaging analysis of selected cells in indicated forebrain assembloids derived from SCZ patients (SCZ #1, 2, 3). (l, n) Quantification of synaptic density, weighted mean firing rate, and spike number per minute of genetically manipulated forebrain assembloids derived from SCZ patients (SCZ #1, 2, 3). (left) Quantification of the synaptic density in indicated forebrain assembloids by calculating the number of synapses per unit area (mm^2^) in each section. Five sections were quantified for one assembloid (Five biological replicates were evaluated; n = 25). Significance was calculated using an unpaired *t*-test. (middle) Quantification of the weighted mean firing rate in the indicated forebrain assembloids measured by MEA. Recordings were performed for thirty minutes. The signals from all active electrodes were quantified for each assembloid (Five biological replicates were evaluated; n = 5). Significance was calculated using an unpaired *t*-test. (right) Quantification of spike number per minute analyzed by calcium imaging of selected cells in indicated forebrain assembloids. The number of spikes was quantified in five selected cells of one assembloid (Five biological replicates were evaluated; n = 25). Significance was calculated using an unpaired *t*-test.

**Supplementary Fig. 9.**
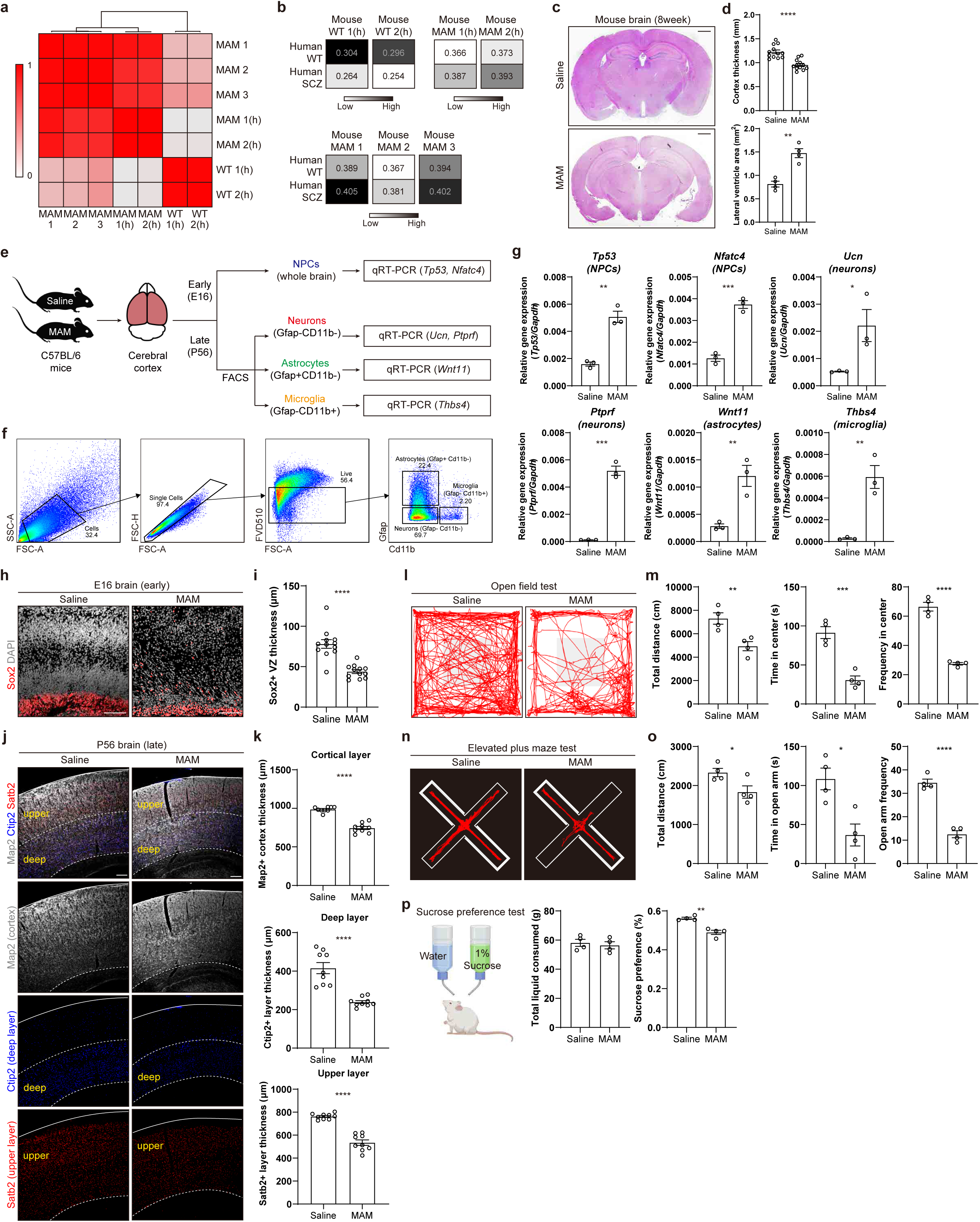
E13 MAM-treated mouse model mimicking brain structural defects and behavior abnormalities of human SCZ. (a) Heat map representation of unsupervised hierarchical clustering from RNA-seq data from wild-type or MAM-treated mouse brain. Samples were clustered based on Pearson correlation coefficients. Samples notated with “(h)” used RNA-seq data from a previous study ^37^. (b) Pearson correlation coefficients obtained through comparing the RNA-seq transcriptome of human wild type or SCZ samples with those of mouse wild-type (top) or MAM (bottom) samples. Samples notated with “(h)” used RNA-seq data from the previous study ^37^. (c) Representative images of coronal sections of the mouse brain treated with saline or MAM analyzed by H&E staining. Scale bars, 1 mm. (d) (top) Quantification of cortical thickness of saline or MAM-treated mice. Three sections per mouse and four mice per group were evaluated (n = 12). Significance was calculated using an unpaired *t*-test. (bottom) Quantification of the lateral ventricle area of saline or MAM-treated mice. Four mice per group were evaluated (n = 4). Significance was calculated using an unpaired *t*-test. (e) Schematic diagram of the gene expression analysis (qRT-PCR) of neural progenitor cells, neurons, astrocytes, and microglia from saline or MAM-treated mice. To analyze the expressions of Tp53 and Nfatc4 in neural progenitor cells, cerebral cortices of saline (control) and MAM-treated mice at E16 were dissociated and qRT-PCR of Tp53 and Nfatc4 was performed. To analyze the expressions of Ucn&Ptprf, Wnt11, and Thbs4 in neurons, astrocytes, and microglia, respectively, cerebral cortices of saline (control) and MAM-treated mice at P56 were dissociated and neurons (Gfap-Cd11b-), astrocytes (Gfap+ Cd11b-), and microglia (Gfap-Cd11b+) were isolated by FACS, which then the qRT-PCR was performed. (f) Gating strategy to analyze neurons (Gfap-Cd11b-), astrocytes (Gfap+ Cd11b-), and microglia (Gfap-Cd11b+). (g) Relative expressions of *Tp53* and *Nfatc4* in neural progenitor cells, *Ucn* and *Ptprf* in neurons, *Wnt11* in astrocytes, and *Thbs4* in microglia isolated from saline or MAM-treated mice following the experimental scheme in Supplementary Fig. 9e. Three biological replicates in each sample were evaluated (n = 3). Significance was calculated using an unpaired *t*-test. (h) Representative images of E16 saline or MAM-treated mouse brain cortex immunostained with Sox2. Scale bars, 50 μm. (i) Quantification of Sox2+ VZ thickness. Three sections per mouse and four mice per group were evaluated (n = 12). Significance was calculated using an unpaired *t*-test. (j) Representative images of P56 saline or MAM-treated mouse brain cortex. Coronal sections were immunostained with Map2, Satb2, and Ctip2. Scale bars, 100 μm. (k) Quantification of the thickness of Map2+ cortical layer (top), Ctip2+ deep layer (middle), and Satb2+ upper layer (bottom) of P56 saline or MAM-treated mouse brain cortex. Three sections per mouse and three mice per group were evaluated (n = 9). Significance was calculated using an unpaired *t*-test. (l) Representative track images of open field tests. (m) Quantification of the total distance traveled (left), time spent in center area (middle), and frequency entering center area (right) of P56 saline or MAM-treated mice. Four mice per group were evaluated (n = 4). Significance was calculated using an unpaired *t*-test. (n) Representative track images for elevated plus maze tests. (o) Quantification of the total distance traveled (left), time spent in open arm (middle), and frequency entering the open arm (right) of P56 saline or MAM-treated mice. Four mice per group were evaluated (n = 4). Significance was calculated using an unpaired *t*-test. (p) A schematic illustration of the two-bottle choice for the sucrose preference test. Quantification of the total amount of liquid consumed (left) and the sucrose preference (right) of P56 saline or MAM-treated mice. Four mice per group were evaluated (n = 4). Significance was calculated using an unpaired *t*-test.

**Supplementary Fig. 10.**
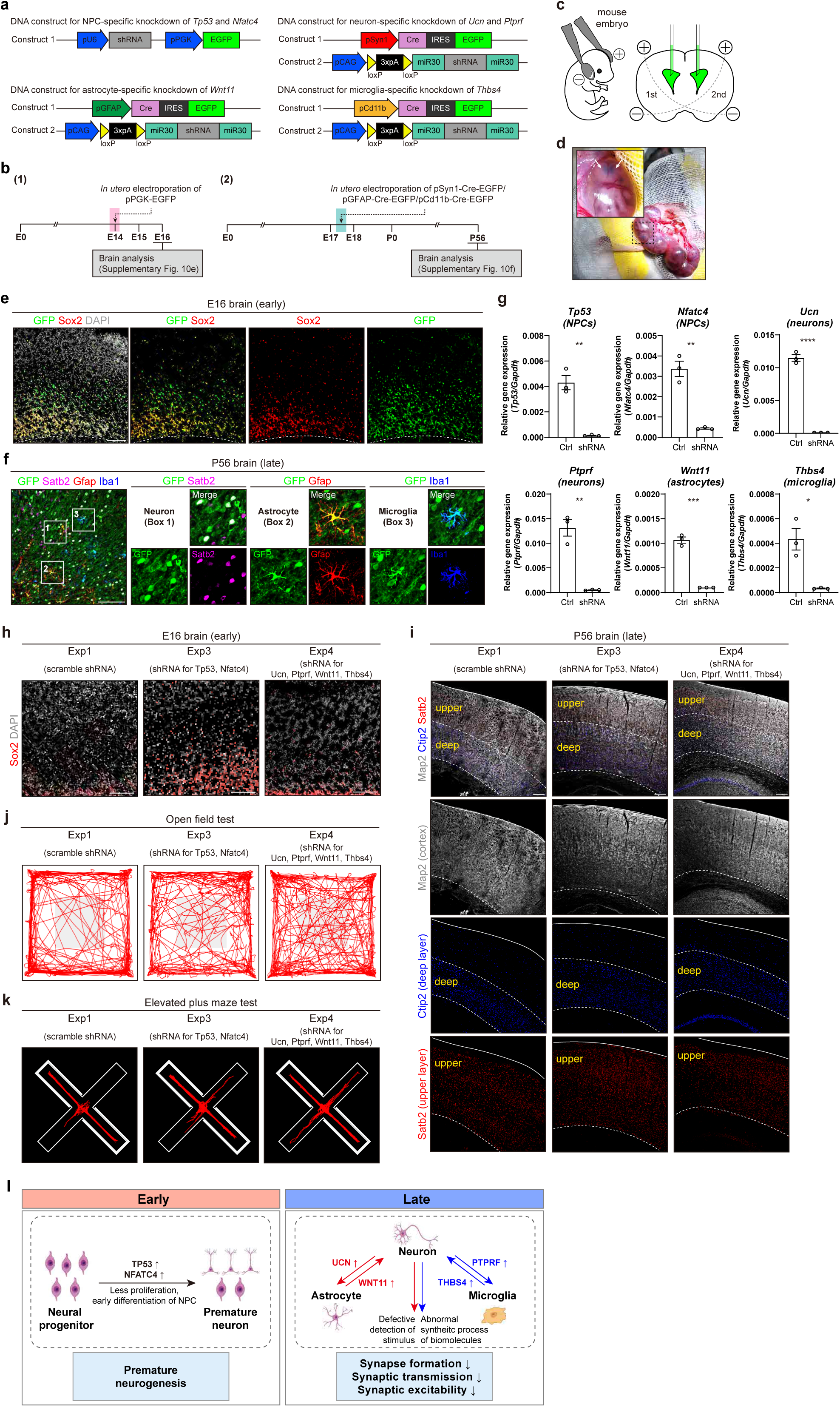
Restoration of structural, functional, and behavioral defects in chemically-induced mouse model of human SCZ through *in utero* genetic manipulations of master regulators. (a) Illustration of DNA constructs used in this study for cell type-specific gene knockdown by *in utero* electroporation. For gene knockdown in neural progenitor cells, electroporation was conducted through using a single DNA construct containing U6 promotor-induced shRNA and PGK promotor-induced EGFP. For cell type-specific gene knockdown in neurons, astrocytes, and microglia, respectively, electroporation was conducted through two DNA constructs each: construct 1 contains Cre and EGFP expression under a cell type-specific promotor; construct 2 contains shRNA whose expression is induced by Cre-mediated recombination. (b) Experimental scheme to validate cell type-specific targeting. To check for NPC-specific targeting, in utero electroporation was conducted in E14 mice with DNA containing pPGK-EGFP (construct 1; upper left in panel ‘a’), followed by brain analysis at E16. To check for neuron-, astrocyte-, and microglia-specific targeting, in utero electroporation was conducted in E17.5 mice with DNA containing pSyn1-Cre-EGFP (construct 1; upper right in panel ‘a’), pGFAP-Cre-EGFP (construct 1; lower left in panel ‘a’), and pCd11b-Cre-EGFP (construct 1; lower right in panel ‘a’), respectively, followed by brain analysis at P56. Targeted cells will exhibit EGFP expression. (c) Schematic representation of the *in utero* electroporation into the mouse brain cortex. (d) Representative images of microinjection into the lateral ventricles of the mouse embryonic brain. Arrows indicate the lateral ventricles filled with DNA-containing solution colored with Fast Green. (e) Representative images of E16 mouse brain cortex electroporated at E14 with DNA containing pPGK-EGFP (construct 1 for NPCs; upper left in panel ‘a’). Coronal sections were immunostained with Sox2. Scale bars, 100 μm. (f) Representative images of P56 mouse brain cortex electroporated at E17.5 with DNA containing pSyn1-Cre-EGFP (construct 1 for neurons; upper right in panel ‘a’), pGFAP-Cre-EGFP (construct 1 for astrocytes; lower left in panel ‘a’), and pCd11b-Cre-EGFP (construct 1 for microglia; lower right in panel ‘a’). Coronal sections were immunostained with Map2, Gfap, or Iba1. Scale bars, 100 μm. (g) Relative expressions of *Tp53* and *Nfatc4* in neural progenitor cells, *Ucn* and *Ptprf* in neurons, *Wnt11* in astrocytes, and *Thbs4* in microglia isolated from MAM-treated mice electroporated with DNA containing scramble (Exp1) or Tp53/Nfatc4/Ucn/Ptprf/Wnt11/Thbs4 shRNA (Exp2) following the experimental scheme in Supplementary Fig. 9e. Three biological replicates in each sample were evaluated (n = 3). Significance was calculated using an unpaired *t*-test. (h) Representative images of E16 MAM-treated mouse brain cortex electroporated with DNA containing scramble (Exp1), Tp53/Nfatc4 shRNA (Exp3) or Ucn/Ptprf/Wnt11/Thbs4 shRNA (Exp4). Coronal sections were immunostained with Sox2. Scale bars, 100 μm. (i) Representative images of P56 MAM-treated mouse brain cortex electroporated with DNA containing scramble (Exp1), Tp53/Nfatc4 shRNA (Exp3) or Ucn/Ptprf/Wnt11/Thbs4 shRNA (Exp4). Coronal sections were immunostained with Map2, Satb2, and Ctip2. Scale bars, 100 μm. (j) Representative track images of open field tests. (k) Representative track images of elevated plus maze tests. (l) A two-step, multifactorial mechanism by which human schizophrenia brains develop. At the early stage, premature neurogenesis leads to the weakening of the laminar organization of the cortical layers, and at later stages, aberrant cellular interactions between neurons and glial cells reduce synapse connectivity.

## Supplementary Data

**Supplementary Data 1. Clinical information of 8 SCZ patients used in this study** Characteristics including age, sex, and race of 8 SCZ patients used in this study are listed. The patients’ cells were purchased from The Coriell Institute (catalog IDs are provided in the chart).

**Supplementary Data 2. Data for transcriptome analysis of early forebrain organoids from healthy individuals and SCZ patients**

DEGs, pathways (from GO analysis), and TFs (from TF enrichment test) are listed. GO analysis and TF enrichment test were performed using DEGs from early forebrain organoids derived from healthy individuals and SCZ patients.

**Supplementary Data 3. Data for transcriptomic analysis of post-mortem SCZ brains**

SCZ DEGs and SCZ-associated modules (from module enrichment test) are listed. The SCZ-associated modules were identified by performing GSEA on previously-reported developmental modules using SCZ DEGs identified from post-mortem brain analysis.

**Supplementary Data 4. Data for transcriptome analysis of genetically-manipulated early forebrain organoids from SCZ patients**

DEGs and pathways (from GO analysis) are listed. GO analysis was performed using DEGs from genetically-manipulated early forebrain organoids derived from SCZ patients.

**Supplementary Data 5. Data for transcriptome analysis of genetically-manipulated early forebrain organoids from healthy individuals**

DEGs and pathways (from GO analysis) are listed. GO analysis was performed using DEGs from genetically-manipulated early forebrain organoids derived from healthy individuals.

**Supplementary Data 6. Data for transcriptome analysis of iPSC-derived and assembloid-derived neurons, astrocytes, and microglia from SCZ patients**

DEGs and pathways (from GO analysis) are listed. GO analysis was performed using DEGs from iPSC-derived and assembloid-derived neurons from SCZ patients.

**Supplementary Data 7. Data for regulatory potential scores of candidate genes expressed in neurons, astrocytes and microglia isolated from SCZ forebrain assembloids**

Data for regulatory potential scores of candidate genes expressed in the neurons, astrocytes, and microglia isolated from SCZ forebrain assembloids, calculated using NicheNet.

**Supplementary Data 8. Data for regulatory potential scores of candidate genes expressed in the neurons, astrocytes and microglia isolated from post-mortem SCZ brains**

Data for regulatory potential scores of candidate genes expressed in the neurons, astrocytes and microglia isolated from post-mortem schizophrenia brains, calculated using NicheNet.

